# Biosynthesis and Physiological Significance of Organ-Specific Flavonol Glycosides in Solanaceae

**DOI:** 10.1101/2025.03.27.645607

**Authors:** Doosan Shin, Haohao Zhao, Ethan Tucker, Keun Ho Cho, Dake Liu, Zhixin Wang, Scott Latimer, Gilles Basset, Yu Wang, Yousong Ding, Jeongim Kim

## Abstract

Flavonols are subclasses of flavonoids, with hundreds of structures identified in plants. This chemical diversity primarily arises from glycosylation, where sugars are selectively added to the flavonol backbone. While flavonol profiles vary across species and organs, the evolutionary forces shaping this chemodiversity and the physiological significance of specific glycosides remain a mystery. Here, we reveal that finely tuned transcriptional regulation and the sugar selectivity of glycosyltransferases drive the formation of distinct organ specific flavonol profiles and a specific flavonol is necessary for male fertility. In Solanaceae pollen, two flavonol glycosides, K2 (kaempferol 3-*O*-glucosyl(1 → 2)galactoside) and Q2 (quercetin 3-*O*-glucosyl(1 → 2)galactoside), are exclusively accumulated. K2 is evolutionarily conserved, while Q2 was lost over time. Consistently, K2 is essential for male fertility, whereas Q2 and aglycones fail to rescue fertility defects. These findings suggest that individual flavonol glycosides have distinct physiological roles, either actively maintained or discarded through evolutionary selection.

## Introduction

Plants produce millions of specialized metabolites, some of which are unique to specific families or species, while others are nearly ubiquitous throughout in the plant kingdom. Flavonoids are a major class of specialized metabolites widely distributed in plants(*1*). The core structure of flavonoids, a C6-C3-C6 skeleton, consists of two aromatic rings (A and B) and a heterocyclic ring (C)(*2*). Based on the structure of the C-ring, flavonoids are classified into subclasses, including flavones, flavonols, flavanones, flavanols, anthocyanidins, and isoflavones(*2*).

Flavonols, in particular, are distinguished by their 3-hydroxyflavone backbone, which can be further modified through hydroxylation, methylation, acylation, and glycosylation resulting in various flavonol structures(*2*). For example, the flavonol aglycones kaempferol and quercetin differ by the presence or absence of a hydroxyl group at the 3′ position of the B-ring, a modification mediated by flavonoid 3′-hydroxylase (F3′H)(*3*, *4*). Among these modifications, glycosylation is the most common and significantly contributes to the remarkable structural diversity of flavonols due to variations in sugar types, attachment positions, and the number of sugar units attached to aglycones(*5*). This glycosylation is primarily catalyzed by UDP-sugar dependent glycosyltransferases (UGTs), which selectively transfer sugar moieties to hydroxyl groups on flavonols(*6*, *7*). These enzymes feature a conserved 44 amino acid PSPG motif in their C-terminal region that is required for glycosylation activity(*7*, *8*). UGTs transfer various UDP-sugars to nucleophilic acceptor sites, typically the hydroxyl groups at positions 3, 5, 7, 3′, or 4′ on flavonol aglycones or on their glycosides, forming inter-glycosidic linkages(*9–14*). This enzymatic versatility enables plants to generate a wide range of glycosidic flavonol structures.

Although flavonols are widely present in plants, the composition and content of flavonol glycosides vary among different plant species and across various organs within a species(*15–19*). In Arabidopsis, for instance, 35 flavonol structures have been identified in various organs, which result from the glycosylation of three aglycones (kaempferol, quercetin, and isorhamnetin) decorated with various sugars such as glucose, rhamnose, and arabinose(*9*). Kaempferol glycosides carrying glucose and rhamnose accumulate ubiquitously in most organs of Arabidopsis, whereas certain flavonols such as isorhamnetin glycosides are restricted to specific organs like flowers(*9*). Quercetin 3-*O*-rhamnosyl(1→6)glucoside (rutin) is commonly found in various monocots and dicots, including buckwheat, citrus trees, apple trees, tomato, and tea plants(*20–24*), but Arabidopsis does not produce rutin, although it accumulates kaempferol 3-*O*-rhamnosyl(1→6)glucoside-7-*O*-rhamnoside(*9*). Despite extensive research on flavonols, it remains largely unknown whether specific flavonol structures are evolutionarily conserved within plant families or in specific organs and how these complex glycosylated flavonols have been shaped and maintained throughout evolution.

Flavonols are generally recognized as antioxidants that contribute to plant stress adaptation(*25*). Multiple studies have also shown that changes in flavonol content or composition influence plant growth and development(*25–31*). For example, the Arabidopsis *F3’H* mutant (*tt7*), which lacks quercetin and its derivatives, exhibits a slower rate of inflorescence growth(*32*). Disruption of flavonol biosynthesis genes such as flavonol synthase (*FLS1*) in Arabidopsis or flavonoid 3-hydroxylase (*F3H*) in tomato causes altered root architectures such as lateral root and root hair development(*33*, *34*). Flavonols are also known to play essential roles in male fertility across various species including tomato, maize, petunia, rice, apple and tobacco(*25*, *28–31*, *35*, *36*). In maize, *chs* mutants lacking all flavonoids showed male sterility, which was rescued with flavonol aglycones, but not with anthocyanidins, flavones or flavanones(*37*).

Two key signaling molecules, reactive oxygen species (ROS) and auxins, are proposed to be involved in flavonol-related plant growth and development phenotypes(*38*, *39*). Plants with reduced flavonol contents showed decreased ROS scavenging capacity, ultimately altering ROS homeostasis and affecting ROS regulated plant development(*38*). Similarly, changes in flavonol content or composition lead to altered auxin transport, a vital process for plant growth regulation(*40*).

Cumulative evidence indicates the pivotal roles of flavonols in plant stress adaptation and plant growth and development. However, any physiological relevance of individual flavonol glycosides remains largely unexplored, although flavonols are collectively recognized as antioxidants. Given the structural diversity of glycosylated flavonols, understanding any specific biological significance of individual flavonol structure is crucial for unraveling their broader role in plant physiology.

In this study, we demonstrate that two flavonols predominantly accumulate in the pollen of various Solanaceae species and one of them, kaempferol 3-O-glucosyl(1 → 2)galactoside (K2), is indispensable for pollen tube growth and successful seed production, while the other pollen-specific flavonol, quercetin 3-O-glucosyl(1 → 2)galactoside (Q2), has been lost over time without functional consequences. This pollen-specific flavonol production is driven by transcriptional regulation and the constrained sugar selectivity of glycosyltransferases. This study also reveals that key amino acid residues dictate the sugar specificity of a pollen-specific flavonol galactosyltransferase.

## Results

### Organ-specific flavonol profiles in tomato

To identify flavonols accumulated in various organs of tomato, we analyzed methanol soluble metabolites prepared with eleven different organs or tissues of the tomato plant (*Solanum lycopersicum*) using High-Performance Liquid Chromatography (HPLC). The analyzed organs included mature pollen grains, leaves, hypocotyls, internodes, roots, petals, sepals, as well as the fruit flesh and peel of both green and red fruits. In all organs except pollen grains, two flavonol rutinosides, quercetin 3-*O*-rhamnosyl(1→6)glucoside (Q-glu-rha, also known as rutin, Q4) and kaempferol 3-*O*-rhamnosyl(1→6)glucoside (K-glu-rha, K4), were predominantly accumulated, with Q4 (rutin) being the most abundant flavonol (Fig. 1, A and B). Among the analyzed organs, red fruit peel showed the highest levels (Fig. 1B, and Fig. S2, and Table S1). However, pollen showed a unique flavonol accumulation pattern. It predominantly accumulated a compound that we called Peak 1, which was below the detection level in all other tested organs except petal (Fig. 1, A and B). To determine the structure of this pollen specific compound, we collected the HPLC eluent corresponding to the retention time of Peak 1 for MS analysis. The content of eluent exhibited an *m/z* value of 611.1594 in positive mode, which corresponds to a calculated exact mass of 610.1521 Da. (Fig. S1A). Further fragmentation analysis of the precursor ion revealed two key fragment ions with *m/z* values of 287.0540 and 465.1009, corresponding to kaempferol and kaempferol-monohexose, respectively (Fig. S1A). The *m/z* differences between the parental ion and two fragments (324.1054 and 146.0512) suggest that Peak 1 represents a kaempferol di-glycoside comprising two hexose units. It was previously shown that the most abundant flavonol glycoside in pollen grains of petunia, a member of the Solanaceae family, is kaempferol 3-*O*-glucosyl(1 → 2)galactoside (K-gal-glu), followed by quercetin 3-*O*-glucosyl(1 → 2)galactoside (Q-gal-glu, hereafter called Q2)(*41*, *42*). We then analyzed the methanolic extract of petunia pollen grains and found that the retention time and UV absorbance spectra of Peak 1 exactly matched those of K-gal-glu, the most abundant peak of the petunia pollen extract (Fig. S1B). Thus, we concluded that Peak 1 is kaempferol 3-*O*-glucosyl(1 → 2)galactoside (K-gal-glu, hereafter called K2)(Fig. 1). This analysis also revealed that Q2 is undetectable in tomato pollen, unlike in petunia pollen (Fig. S1B).

**Fig. 1.**
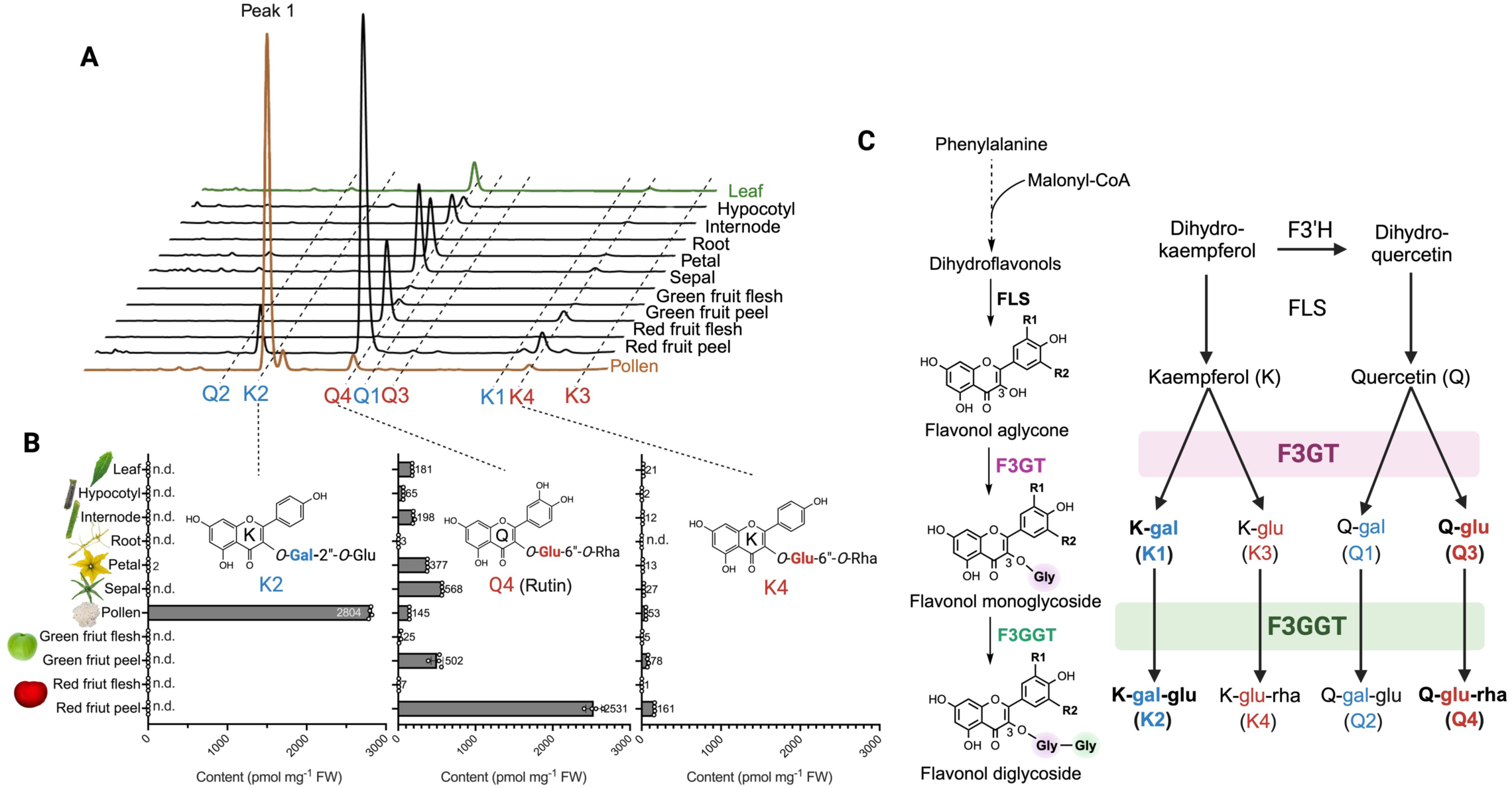
Flavonol profiling with eleven tomato organs and tissues reveals pollen-specific compounds. **(A)** Representative HPLC chromatograms of methanolic extracts from eleven tomato organs and tissues, together with authentic standards of four 3-*O*-glucosylated flavonols (K3, K4, Q3, and Q4) and two flavonol 3-*O*-galactosides (K1 and Q1). The pollen extract shown in the chromatogram was extracted in 50% methanol at a concentration approximately 6.2 times more diluted compared to other organs. Peak 1 indicated with a blue dot is highly accumulated in pollen grains, while no peak 1 was detected in the rest of samples. Peak 1 characterized as K2 (kaempferol 3-*O*-glucosyl(1→2)galactoside) in Figure S1. Chlorogenic acid, a caffoylquinate, are marked in the chromatograms. **(B)** Quantification of three major flavonol glycosides (K2, K4 and Q4) across different tomato organs detected with HPLC. The data presented are the mean concentrations (pmol mg⁻¹ fresh weight) of flavonol glycosides and their standard deviations (n = 4). Compounds that were below the detection limit in certain organs are marked as not detected (n.d.). **(C)** Simplified biosynthetic pathway of flavonol glycosides in tomato. Flavonol biosynthesis begins with the conversion of phenylalanine into dihydroflavonols, mediated by several enzymatic steps. Dihydroflavonols are then converted to flavonols (kaempferol and quercetin) by flavonol synthase (FLS). Kaempferol (K) and quercetin (Q) undergo glycosylation by flavonol 3-*O*-glycosyltransferases (F3GT), leading to the production of initial glycosylated intermediates, such as kaempferol 3-*O*-galactoside (K-gal; K1), kaempferol 3-*O*-glucoside (K-glu; K3), quercetin 3-*O*-galactoside (Q-gal; Q1), and quercetin 3-*O*-glucoside (Q-glu; Q3). These intermediates are further modified by flavonol 3-*O*-glycoside: glycosyltransferases (F3GGT), generating secondary glycosides, including kaempferol 3-*O*-glucosyl(1→2)galactoside (K-gal-glu; K2), kaempferol 3-*O*-glucosyl(1→6)rhamnoside (K-glu-rha; K4), quercetin 3-*O*-glucosyl(1→2)galactoside (Q-gal-glu; Q2), and quercetin 3-*O*-glucosyl(1→6)rhamnoside (Q-glu-rha; Q4). Abbreviations used in chemical structures are K, kaempferol; Q, quercetin; gal, galactose; glu, glucose; rha, rhamnose. Kaempferol has hydrogen (H) at both R1 and R2, whereas quercetin has H at R1 and a hydroxyl group (OH) at R2.

To detect low abundant flavonols, we reanalyzed the same methanolic soluble extracts with LC-MS and detected eight flavonols that matched authentic standards: kaempferol (K), quercetin (Q), K-glu (K3), Q-glu (Q3), Q-gal (Q1), K-gal (K1), K-glu-rha (K4) and Q-glu-rha (Q4) (Fig. 1C). The compound having the exact mass of 610.1534 Da was again detected in pollen extracts, which matches the mass of the K2 compound (Fig. S2, and Table S1). Flavonol rutinosides (Q4 and K4) accumulated across various organs and their mono-glucoside precursors, K-glu (K3) and Q-glu (Q3), were detected in K4 and Q4 enriched organs (Fig. 1B, and Fig. S2, and Table S1). In contrast, K2 (K-gal-glu) and K1 (K-gal) accumulated predominantly in pollen and two additional flavonol triglycosides (FGGG-1 and FGGG-2) were also detected in various organs (Fig. S2, and Table S1).

### Identification of the enzymes responsible for the production of organ-specific flavonol glycosides

Detected flavonol glycosides contain either glucose or galactose at the 3-OH position of the flavonol aglycones, kaempferol (K) or quercetin (Q) (Fig. 1, and Fig. S2), which requires flavonol 3-*O*-glycosyltransferase (F3GT) activity (Fig. 1C). The production of flavonol 3-*O*-diglycosides, such as K-gal-glu (K2), requires flavonol 3-*O*-glycoside: glycosyltransferase (F3GGT) activity (Fig. 1C). To identify the UGTs with F3GT or F3GGT activities, we conducted a phylogenetic analysis using protein sequences of 182 putative UGTs identified from the tomato genome (SL4.0 build; ITAG4.0 annotation)(*43*). Our analysis also included 13 previously characterized F3GTs and F3GGTs from Arabidopsis, petunia, eggplant, tea, grape, barley, and maize(*6*, *44–53*). Two UGT homologs, Solyc10g083440 and Solyc07g006720, were clustered with other characterized F3GTs in the F3GT clade (Fig. 2A). It was previously shown that Solyc10g083440 (SlUGT78D-A) can convert quercetin to quercetin 3-*O*-glucoside (*24*), but the function of Solyc07g006720 (hereafter called SlUGT78D-B) remains unknown. According to the public database, *SlUGT78D-A* is expressed in most organs except pollen and root, whereas *SlUGT78D-B* is highly expressed in pollen, which we confirmed experimentally (Fig. 2, B and C).

**Fig. 2.**
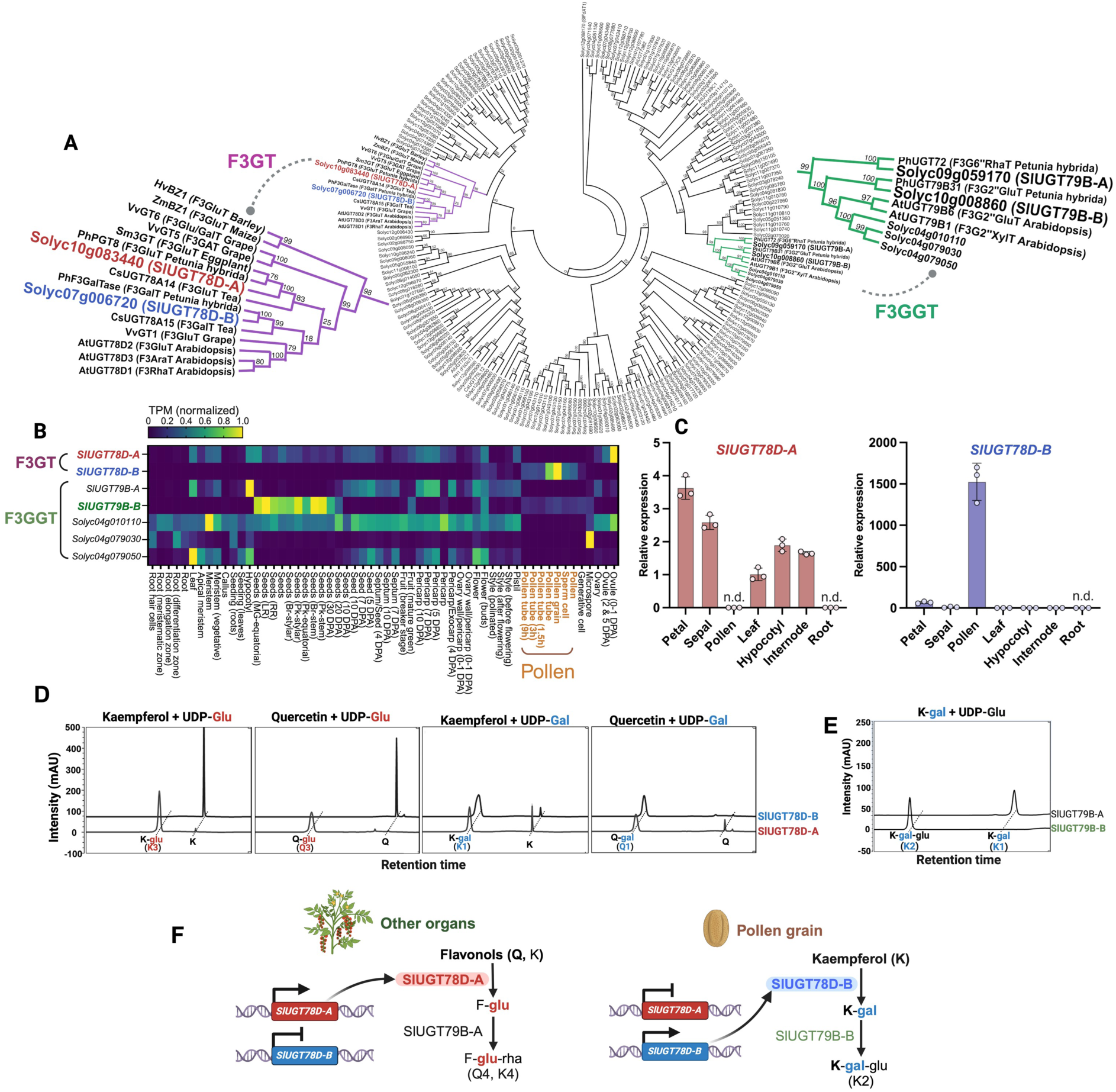
Identification of two UGT enzymes responsible for a pollen-specific flavonol glycosides. **(A)** A phylogenetic tree of UDP-Glycosyltransferases (UGTs). A phylogenetic tree constructed using 182 UGT protein sequences retrieved from the tomato genome and previously characterized F3GTs and F3GGTs from other species (Table S4). The tree was generated using the maximum likelihood method with 1000 bootstrap replicates and WAG with Freqs. (+F) model. Solyc12g088170 (anthocyanin acyltransferase; SlFdAT1) was used as an outgroup. Two zoom-in clades representing the F3GT and the F3GGT clades are shown in purple and green, respectively. **(B)** Heatmap showing the normalized expression (TPM) of *UGT* genes in different tomato organs, including roots, stems, leaves, flowers, and fruits. Particularly, *SlUGT78D-B* and *SlUGT79B-B* show expression in pollen while other flavonol UGT homologs do not. The expression data were sourced from a public database (https://conekt.sbs.ntu.edu.sg)(91) and raw expression data are available in Table S2. **(C)** The relative gene expression levels of *SlUGT78D-A* and *SlUGT78D-B* in different tomato organs, measured by qRT-PCR. Expression was normalized to *SlPP2Acs* (*Solyc05g006590*) an internal control. The data are presented as mean ± SD (n = 3). n.d. indicates not detected **(D)** In vitro enzymatic assays of SlUGT78D-A and SlUGT78D-B. HPLC chromatograms of enzyme reaction mixtures demonstrate the production of K-glu (K3), Q-glu (Q3), K-gal (K1), and Q-gal (Q1) when SlUGT78D-A and SlUGT78D-B were incubated with two flavonol aglycones, kaempferol (K) and quercetin (Q), and two sugar donors, UDP-glucose and UDP-galactose. SlUGT78D-A shows both glucosyltransferase and galactosyltransferase activities at the 3-OH position on kaempferol and quercetin, while SlUGT78D-B exhibits galactosyltransferase activity, but no glucosyltransferase activity. Chromatograms were obtained at a wavelength of 345 nm. **(E)** HPLC chromatograms showing the enzymatic activity of SlUGT79B-B, which produces K-gal-glu (K2) from K-gal using UDP-glucose as sugar donor. SlUGT79B-A does not show this activity. The chromatograms were obtained at a wavelength of 345 nm. **(F)** Working model of transcriptional regulation of key enzymes determining the distinct flavonol glycoside profiles between pollen and other organs. In organs outside of pollen, flavonoid 3’-hydroxylase (F3’H) is active, producing both kaempferol and quercetin, which are glucosylated by SlUGT78D-A to form flavonol 3-*O*-glucosides (F-glu). Further glycosylation by SlUGT79B-A results in the formation of flavonol 3-*O*-glucosyl(1→6)rhamnoside (F-glu-rha; Q4, K4). In pollen, however, F3’H expression is repressed, and galactosylation becomes predominant due to the activation of SlUGT78D-B and repression of SlUGT78D-A. This switch leads to the production of kaempferol 3-*O*-galactoside (K-gal). Further modification by SlUGT79B-B results in kaempferol 3-*O*-glucosyl(1→2)glucoside (K2), which is uniquely abundant in pollen. Abbreviations are K, kaempferol; Q, quercetin; UDP-Glu, UDP-glucose; UDP-Gal, UDP-galactose; glu, glucose; gal, galactose; rha, rhamnose.

Then, we conducted enzyme activity tests using recombinant SlUGT78D-A and SlUGT78D-B with two sugar donors, UDP-glucose and UDP-galactose, and two sugar acceptors, kaempferol and quercetin. SlUGT78D-A produced flavonol 3-*O*-glucosides (K3 and Q3) or flavonol 3-*O*-galactosides (K1 and Q1) (Fig. 2D), indicating that it has both flavonol 3-*O*-glucosyltransferase (F3Glu) and 3-*O*-galactosyltransferase (F3Gal) activities. In contrast, SlUGT78D-B produced flavonol 3-*O*-galactosides, but not flavonol 3-*O*-glucosides, suggesting that SlUGT78D-B functions as a flavonol 3-*O*-galactosyltransferase (F3Gal) but lacks 3-*O*-glucosyltransferase activity (F3Glu) (Fig. 2D). It is noteworthy that flavonol 3-*O*-diglucosides, K4 and Q4, were detected in pollen grains, but SlUGT78D-A having F3Glu activity is not expressed in pollen grains and the pollen-specific F3GT, SlUGT78D-B, does not possess F3Glu activity in vitro (Fig. 2, B and C). It appears that additional enzymes with F3Glu activity exist in pollen or all 3-*O*-glucosylated flavonols such as Q4 are transported from other organs or tissues like tapetum(*54*). However, our phylogenetic analysis did not identify any additional UGTs other than SlUGT78D-A and SlUGT78D-B within the F3GT clade (Fig. 2A).

The production of K-gal-glu (K2) requires an additional UGT enzyme that can add glucose to K-gal at its 2’’ position (2’’-*O*-glucosyltransferase activity), which requires flavonol 3-*O*-glycoside: glycosyltransferase (F3GGT) activity (Fig. 1C). In the F3GGT clade, five putative tomato UGTs were identified (Fig. 2A). Solyc09g059170 (SlUGT79B-A) clustered closely with Petunia flavonoid 3-*O*-glucoside: 6’’-*O*-rhamnosyltransferase, while Solyc10g008860 (SlUGT79B-B) clustered with petunia flavonol 3-*O*-glycoside: 2’’-*O*-glucosyltransferase. The other three enzymes grouped with Arabidopsis F3GGTs (Fig. 2A). However, only *SlUGT79B-B* is expressed in pollen grains, while the other four candidates show no expression in pollen grains according to a public database(*55*) (Fig. 2B, and Table S2). Thus, we hypothesized that SlUGT79B-B is the F3GGT responsible for K2 production in pollen grains. Indeed, recombinant SlUGT79B-B showed kaempferol 3-*O*-glycoside: 2’’-*O*-glucosyltransferase activity as it produced K-gal-glu (K2) using K-gal and UDP-Glucose, while recombinant SlUGT79B-A did not show the same activity (Fig. 2E). Our results indicate that differentially expressed flavonol glycosyltransferases and their sugar specificities shape the distinctive organ-specific flavonol glycoside profiles in tomato. In pollen, the repression of *SlUGT78D-A* together with the activation of *SlUGT78D-B* enriches kaempferol galactosides, while the predominant activity of *SlUGT78D-A* with repressed *SlUGT78D-B* results in abundant flavonol glucosides in other organs (Fig. 2G).

### Key residues determining sugar specificity of SlUGT78D-B

SlUGT78D-A and SlUGT78D-B showed not only unique organ-specific expression patterns but also distinct sugar specificities, despite both enzymes having relaxed substrate specificities toward flavonol aglycones (Fig. 2D). To determine the molecular basis of their differing sugar preferences, we compared the amino acid sequences of SlUGT78D-A and SlUGT78D-B homologs. Our analysis included the characterized CsUGT78A14 (flavonol 3-*O*-glucosyltransferase, F3GluT) and CsUGT78A15 (flavonol 3-*O*-galactosyltransferase, F3GalT) from *Camellia sinensis*(*51*) and five Solanaceae species (identified via an NCBI GenBank BLAST search) (Fig. 3A, and Fig. S4). Phylogenetic analysis grouped the homologs into two distinct clades: the F3GluT group (including SlUGT78D-A) and the F3GalT group (including SlUGT78D-B) (Fig. 3A). The homologs within each group shared over 75% sequence similarity, while similarity between groups was less than 50% (Fig. S4). We identified 11 amino acids conserved in each group (Fig. S5), and also identified 17 residues predicted to be exposed in the ligand-binding pocket based on studies of GT1 from *Vitis vinifera*(*6*, *50*) (Fig. S5), leading us to pinpoint four residues in SlUGT78D-B (14A, 19L, 138S, and 373H) which are positioned near the predicted sugar binding site (Fig. 3, B and C, and Fig. S5).

**Fig. 3.**
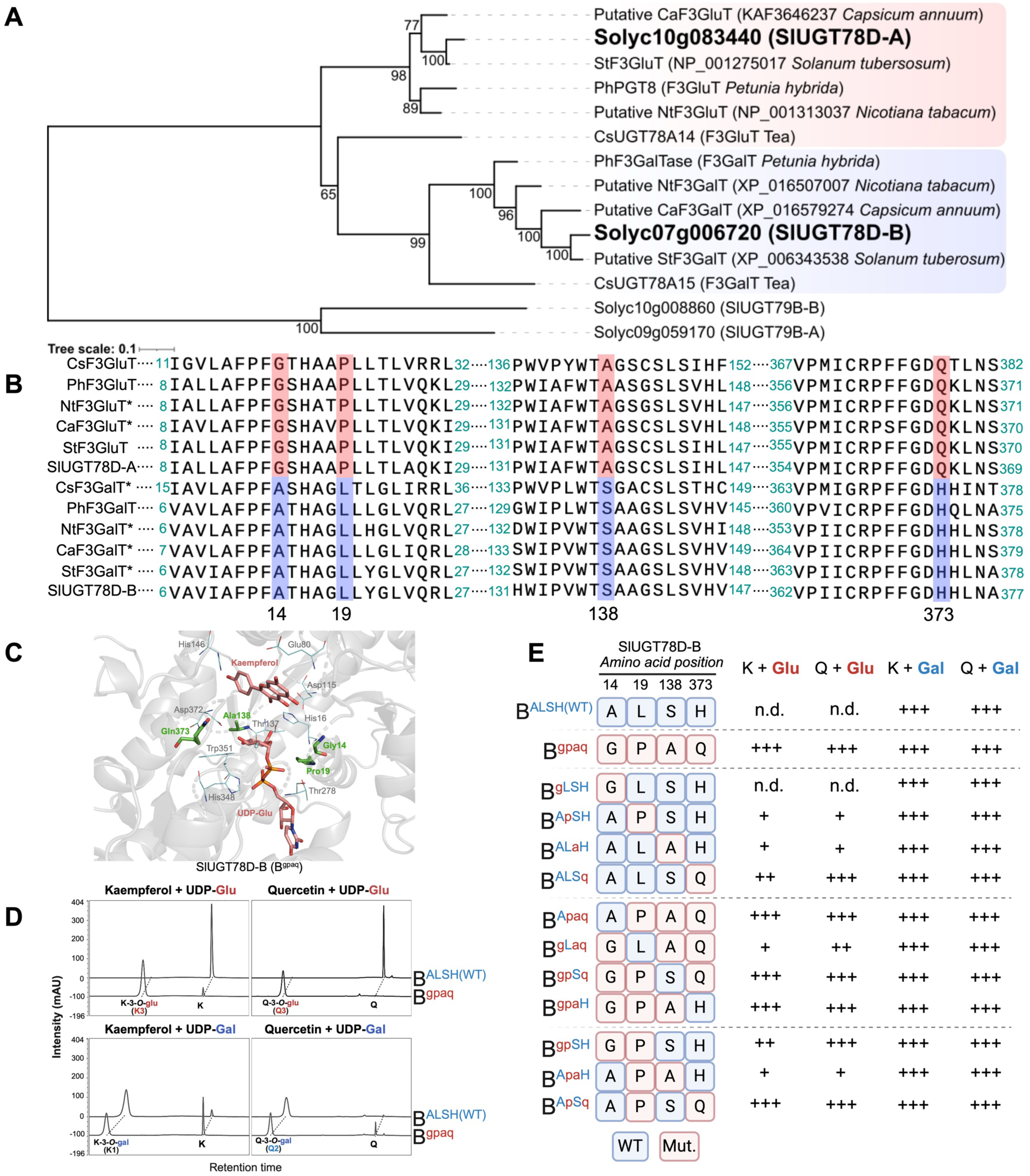
Identification of key amino acids for sugar specificity in SlUGT78D-B. **(A)** Phylogenetic analysis with SlUGT78D-A and SlUGT78D-B homologs from Solanaceae species shows two distinct groups, the SlUGT78D-A group and the SlUGT78D-B group, which are highlighted in red and blue, respectively. The tree was constructed using the maximum likelihood method with a bootstrap analysis of 1000 replicates, using the WAG with Freqs. (+F) model. Tomato F3GGT enzymes, SlUGT79B-B and SlUGT79B-A, are used as the outgroup. **(B)** Sequence alignment of F3GTs showing four amino acid residues conserved in each group, which are highlighted in red and blue for the SlUGT78D-A group and the SlUGT78D-B group, respectively. Amino acids positions 14, 19, 138, and 373 are indicated based on the SlUGT78D-B sequences. **(C)** A structural model of SlUGT78D-B was generated with ColabFold(*91*) and visualized with PyMOL2.5. The ligand position was determined according to the VvGT1 structure (PDB DOI: https://doi.org/10.2210/pdb2C1Z/pdb). Amino acids targeted for point mutations are highlighted in green. **(D)** HPLC chromatograms showing the activity of wild-type SlUGT78D-B (B^ALSH^) and mutated SlUGT78D-B (B^gpaq^) with kaempferol and quercetin substrates, using UDP-galactose and UDP-glucose as sugar donors. The wild-type enzyme (B^ALSH^) shows galactosyltransferase activity, but no glucosyltransferase activity, while the quadruple mutant (B^gpaq^) acquires glucosyltransferase activity, producing K-3-*O*-glu and Q-3*-O*-glu. The HPLC chromatograms were obtained at a wavelength of 345 nm. **(E)** The glucosyltransferase and galactosyltransferase activities of SlUGT78D-B mutant variants. Activities are shown for quadruple, triple, double, and single mutants at positions 14, 19, 138, and 373. The presence of activity is indicated with multiple (+) marks based on product peak area in HPLC analysis (a higher number of ‘+’ marks indicate a greater level of the detected product compound), and the absence of activity is indicated with “n.d.” (not detected).

To test the significance of the identified four residues in sugar specificity, we generated various modified SlUGT78D-B proteins with substitutions at these positions using the corresponding amino acids from SlUGT78D-A. SlUGT78D-B^gpaq^ carrying all four amino acid substitutions (A14G, L19P, S138A, H373Q) gained flavonol 3-*O*-glucosyltransferase activity while retaining its galactosyltransferase activity, thereby resembling the bifunctional activity of SlUGT78D-A (Fig. 3, D and E, and Fig. S6). This suggests that these residues are essential for sugar specificity individually or in combination. To further narrow down key residues, we generated additional modified SlUGT78D-B with single, double, and triple amino acid substitutions and they all exhibited bifunctional activity, except for the single substitution at A14 (Fig. 3E, and Fig. S6). Given that any single amino acid substitution of L19, S138, and H373 enabled SlUGT78D-B to acquire glucosyltransferase activity, there is a selective pressure to restrict its glucosyltransferase activity while maintaining its galactosyltransferase activity.

Notably, introducing the corresponding residues from SlUGT78D-B into SlUGT78D-A abolished both glucosyltransferase and galactosyltransferase activities (Fig. S6), further confirming their critical role for its enzymatic activities.

### Disruption of SlUGT78D-B leads to impaired pollen tube growth and reduced seed yield

To determine any physiological functions of flavonol glycosides in tomato, we disrupted SlUGT78D-A and SlUGT78D-B using a CRISPR/Cas9 system that simultaneously targets both genes. Three independent homozygous mutant lines (named *Slugt78d-a*, *Slugt78d-b*, and *Slugt78d-a/b*) were established (Fig. 4A, and Fig. S7). The *Slugt78d-a* mutant has a 4 bp deletion mutation in the exon of *SlUGT78D-A*, resulting in an early stop codon, while *SlUGT78D-B* remained unaltered. The *Slugt78d-b* mutant has a 2 bp deletion in the first exon of *SlUGT78D-B* that causes an early stop codon. It also carries a 9 bp deletion in *SlUGT78D-A* that removes three amino acids. Finally, the *Slugt78d-a/b* mutant has a 4 bp deletion in the exon of *SlUGT78D-A* and a 22 bp deletion in the first exon of *SlUGT78D-B*, both resulting in early stop codons. All frameshift mutations result in truncated proteins lacking the Plant Secondary Product Glycosyltransferase (PSPG) motif, which is required for the glycosyltransferase reaction(*56*) (Fig. S7).

**Fig. 4.**
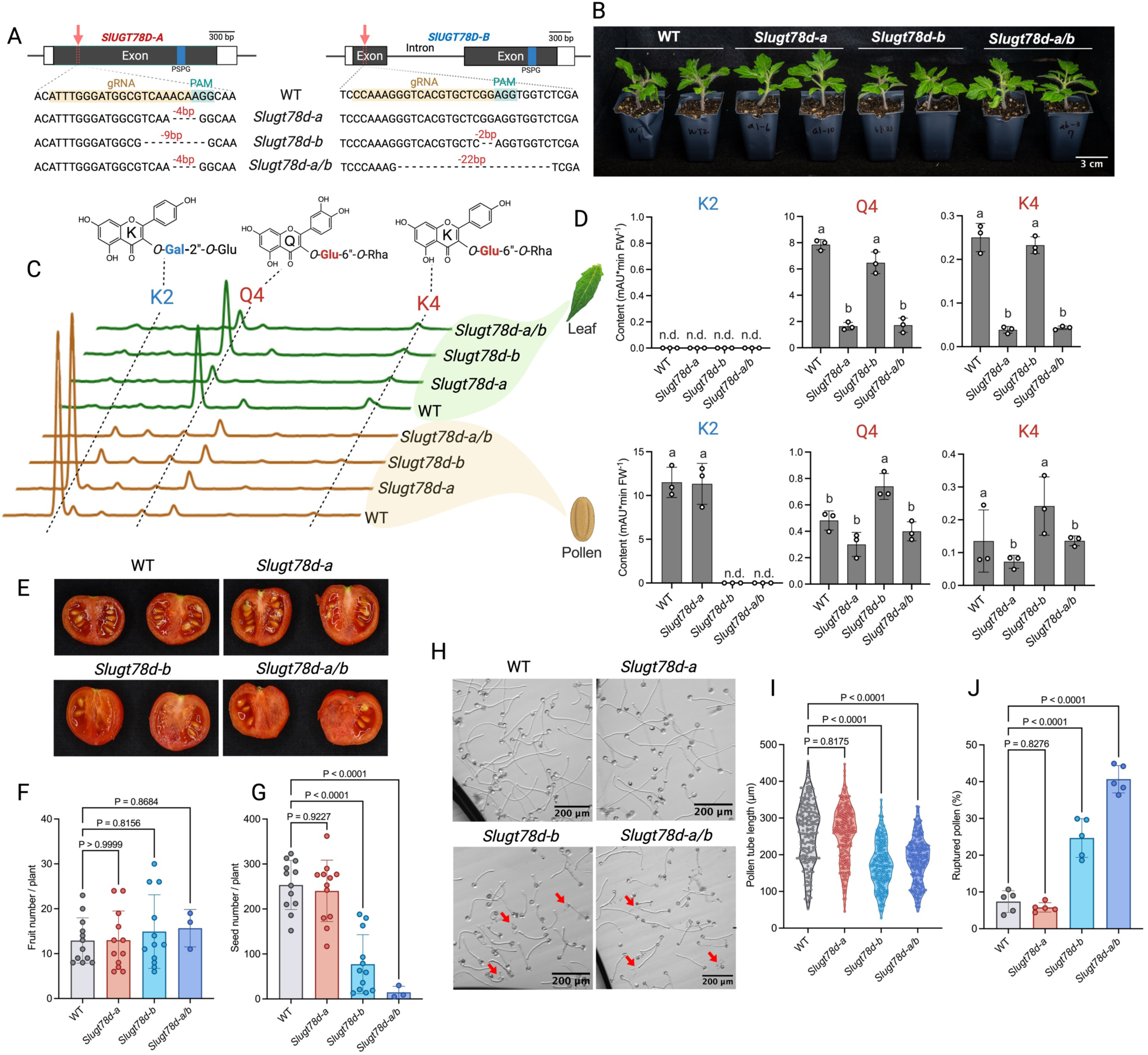
Disruption of SlUGT78D-B abolishes pollen kaempferol galactosylation, leading to defective pollen tube development and lower seed yield. **(A)** Schematic gene structures of *SlUGT78D-A* and *SlUGT78D-B* with the CRISPR-Cas9 targeting sites and mutations in *Slugt78* mutants. PSPG motif is marked on gene structures with a blue box. The guide RNA (gRNA) binding sites and the protospacer adjacent motif (PAM) are highlighted in yellow and green, respectively. The resulting deletion mutations in the mutant lines (*Slugt78d-a*, *Slugt78d-b*, and *Slugt78d-a/b*) are indicated. **(B)** Representative images of 3-week-old wild type (WT), *Slugt78d-a*, *Slugt78d-b*, and *Slugt78d-a/b*. Scale bar = 3 cm. **(C)** Representative HPLC chromatogram of methanolic extracts from leaves (green traces) and pollen (brown traces) of 4-week-old wild type (WT), *Slugt78d-a*, *Slugt78d-b*, and *Slugt78d-a/b* mutants. The retention times for kaempferol 3-O-glucosyl(1→2)galactoside (K2), quercetin 3-O-glucosyl(1→6)rhamnoside (Q4), and kaempferol 3-O-glucosyl(1→6)rhamnoside (K4) are indicated on the chromatogram. The compound structure at the top of the graph are identified by their abbreviations: K for kaempferol, Q for quercetin, glu for glucose, gal for galactose, and rha for rhamnose. **(D)** The levels of K2, Q4 and K4 in leaves and pollen from wild type (WT) and three mutant lines. Methanolic extracts were analyzed with HPLC, and the concentration is shown in peak area per fresh weight (mAU*min FW⁻¹). Data represent mean ± SD (n=3). Statistical significance was determined using one-way ANOVA, followed by Tukey’s post-hoc test, with different letters indicating significant differences among groups (P-value < 0.05). Each data point is shown as an open circle on the bar graph. n.d. indicates not detected. **(E)** Cross-sectioned fruits of WT, *Slugt78d-a*, *Slugt78d-b*, and *Slugt78d-a/b*. **(F,G)** Bar graph showing the average number of fruits per plant (f), and the number of seeds per plant (g) Data represent mean ± SD (n=12 for all groups except *Slugt78d-a/b*, where n=3). **(H)** Microscopic images of growing pollen tubes in WT, *Slugt78d-a*, *Slugt78d-b*, and *Slugt78d-a/b*. Red arrows indicate ruptured pollen tubes. Scale bar = 200 μm. **(I)** Violin plot showing pollen tube length for WT, *Slugt78d-a*, *Slugt78d-b*, and *Slugt78d-a/b* (n = 200)**. (J)** Bar graph showing the percentage of ruptured pollens. Data represent mean ± SD (n=5 with each replicate containing approximately 100 pollen grains)**. (F-G, I-J)** Each data point is shown as a circle. Statistical significance was determined using one-way ANOVA, followed by Dunnett’s multiple comparisons test, with adjusted P-values marked at the top.

None of the mutants exhibited any visible changes in vegetative growth or development (Fig. 4B). However, both *Slugt78d-a* and *Slugt78d-a/b* mutants showed an 80% reduction in the levels of flavonol glucosides, K-glu-rha (K4) and Q-glu-rha (Q4) in their leaves compared to wild type and *Slugt78d-b* mutant (Fig. 4, C and D), which aligns with the flavonol 3-*O*-glucosyltransferase activity of SlUGT78D-A. In contrast, the levels of K4 and Q4 in the pollen of these mutants were not substantially changed compared to wild type (Fig. 4, C and D). The *Slugt78d-b* mutant carries a 9 bp deletion in *SlUGT78D-A*. However, the levels of K4 and Q4 remained unchanged in its leaves, suggesting that the three amino acid deletion does not affect its glucosyltransferase activity (Fig. 4, A, C and D). Thus, we consider *Slugt78d-b* to be a single null mutant of *SlUGT78D-B*.

The pollen-specific flavonol, K-gal-glu (K2), was not detected in both *Slugt78d-b* and *Slugt78d-a/b* pollen, whereas *Slugt78d-a* mutant accumulated wild-type levels of K2 (Fig. 4, C and D) in its pollen. *SlUGT78D-A* expression was not induced in the *Slugt78d-b* pollen (Fig. S8), suggesting that SlUGT78D-B is the sole enzyme responsible for the 3-*O*-galactosylation of kaempferol in pollen.

Fruit shape and the total number of fruits per plant remained unaltered in these mutants compared to wild type (Fig. 4, E and F). However, the number of seeds per plant was significantly reduced in *Slugt78d-b* and *Slugt78d-a/b* compared to wild type and *Slugt78d-a* (Fig. 4G). As successful seed production requires effective fertilization, and flavonoid deficient mutants in several species have been reported to display defective pollen(*29*, *30*, *57–59*), we thought that the reduced seed yield might be related to pollen development. Interestingly, while the overall shape and size of pollen grains are indistinguishable across all genotypes (Fig. S9), the pollen tube lengths in *Slugt78d-b* and *Slugt78d-a/b* were significantly shorter than those in wild type and *Slugt78d-a* (Fig. 4, H and I). The number of ruptured pollen tubes was also increased in *Slugt78d-b* and *Slugt78d-a/b* (Fig. 4, H and J). Both *Slugt78d-b* and *Slugt78d-a/b* mutants lack the pollen-specific kaempferol galactoside (K2) due to the absence of functional SlUGT78D-B. It was previously shown that the tomato flavonol deficient mutant showed deformed pollen grain shape and abnormal pollen tube growth, accompanied with increased levels of reactive oxygen species (ROS) and the supplementation of kaempferol reduced the levels of ROS and rescued defective pollen(*59*, *60*). However, we found that steady state levels of free kaempferol in *Slugt78d-b* and *Slugt78d-a/b* mutants were not reduced compared to the wild type (Fig. S10B). In fact, *Slugt78d-a/b* mutants accumulated a sevenfold higher level of free kaempferol compared to wild type. Kaempferol aglycone is a precursor for ubiquinone, a co-factor of the cellular respiration(*61–63*) (Fig. S10C). The blocked glycosylation of kaempferol in the *Slugt78d-b* mutant may affect the production of ubiquinone, a lipophilic antioxidant.

However, the levels of ubiquinone Q10 were similar in all four genotypes (Fig. S10C).

Since kaempferol aglycone is not limited in *Slugt78d-b* pollen, we hypothesized that the presence of kaempferol 3-*O*-galactoside or its derivatives such as K2 is associated with proper pollen tube growth.

### Kaempferol galactosides, but neither aglycones nor quercetin galactosides, rescued defective pollen tube growth

Since the pollen-specific galactoside, K-gal-glu (K2), was detected in the pollen of both tomato and petunia(*41*, *42*) and homologs of the K2 biosynthetic enzymes, SlUGT78D-B and SlUGT79B-B, are found in other Solanaceae plants (Fig. 3A, and Fig. S4, and S11), we thought that K2 might be conserved across other Solanaceae species. To test it, we analyzed methanolic extracts of pollen from *Solanum tuberosum* (potato), *Capsicum annuum* (chili pepper), and two tobacco species (*Nicotiana tabacum* and *Nicotiana benthamiana*) and found that K2 is the most abundant flavonol in all samples (Fig. 5A, and Fig. S12). Interestingly, Q-gal-glu (Q2) was detected in petunia and two tobacco species, but not in chili pepper, potato, and tomato (Fig. 5A). Considering that the chili pepper, potato, and tomato diverged more recently than tobacco and petunia(*64*) (Fig. 5B), K2 appears to have been maintained during the evolution, whereas Q2 was lost over time. We showed that SlUGT78D-B can convert quercetin to quercetin 3-*O*-galactoside, which can be further modified by SlUGT79B-B to produce Q2 (Fig. S1B, and S11). However, tomato pollen does not accumulate Q2. One possible limiting factor is the availability of quercetin in tomato pollen. F3’H activity is required for quercetin production (Fig. 1C), but *SlF3’H* is not expressed in tomato pollen although it expresses in most other organs (Fig. S3).

**Fig. 5.**
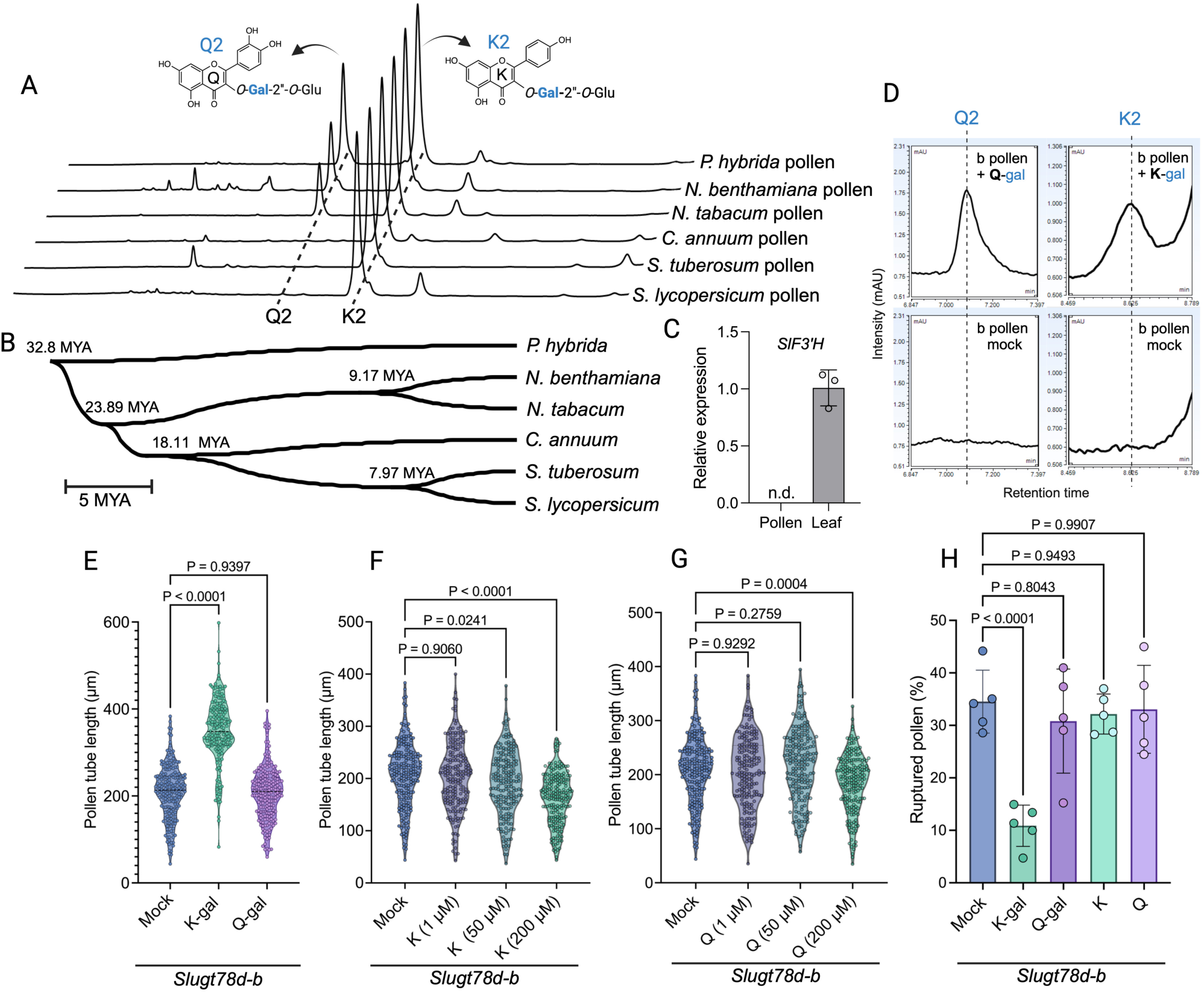
Selective rescue of pollen tube defects by galactosylated kaempferol and its evolutionary implications for galactosylated quercetin loss in late-diverged Solanaceae. **(A)** HPLC chromatograms showing the presence of quercetin 3-*O*-glucosyl(1→2)galactoside (Q2) and kaempferol 3-*O*-glucosyl(1→2)galactoside (K2) in pollen extracts from six Solanaceae plants including *Petunia hybrida* (*P. hybrida*), *Nicotiana benthamiana* (*N. benthamiana*), *Nicotiana tabacum* (*N. tabacum*), *Capsicum annuum* (*C. annuum*), *Solanum tuberosum* (*S. tuberosum*), and *Solanum lycopersicum* (*S. lycopersicum*). The K2 peak was detected in methanolic extracts of all six Solanaceae pollens, while Q2 was detected only in petunia and two tobacco species. The HPLC chromatograms were obtained at a wavelength of 345 nm. **(B)** Phylogenetic tree depicting speciation within the Solanaceae family. The tree illustrates the divergence times (in millions of years ago, MYA) among Solanaceae species shown in (a). Divergence times are labeled at each node. The phylogenetic tree was retrieved from TimeTree, a web-based database (https://timetree.org). The scale bar represents 5 million years (MYA). **(C)** The relative gene expression levels of *SlF3’H* in pollen and leaf, measured by qRT-PCR. Expression was normalized to *SlPP2Acs* (*Solyc05g006590*) an internal control. The data are presented as mean ± SD (n = 3). n.d. indicates not detected**. (D)** Conversion of flavonol monoglycosides to flavonol diglycosides in *Slugt78d-b* mutant pollen in chemical complementation. HPLC chromatograms show the conversion of kaempferol 3-*O*-galactoside (K-gal) to kaempferol 3-*O*-glucosyl(1→2)galactoside (K2) in the left panel, and quercetin 3-*O*-galactoside (Q-gal) to quercetin 3-*O*-glucosyl(1→2)galactoside (Q2) in the right panel, in *Slugt78d-b* pollen treated with 200 μM flavonol monoglycosides. Pollen treated with the same volume of DMSO (mock control) did not produce detectable levels of Q2 or K2. After flavonol monoglycoside treatment in PGM media, metabolites were extracted using a methanolic solution following 4 hours of incubation. Chromatograms were extracted at a wavelength of 345 nm, confirming the formation of flavonol diglycosides. **(E)** Violin plot showing the pollen tube length (n = 200) of *Slugt78d-b* pollen treated with 200 μM flavonol monoglycosides (kaempferol 3-*O*-galactoside, K-gal; and quercetin 3-*O*-galactoside, Q-gal) or an equivalent volume of DMSO (mock). **(F, G)** Violin plots showing the pollen tube length of *Slugt78d-b* pollen treated with varying concentrations (1 μM, 50 μM, and 200 μM) of kaempferol (K) and quercetin (Q) aglycones (n = 200). **(H)** Bar graph illustrating the percentage of ruptured pollen tubes (n = 5, with each replicate containing approximately 100 pollen grains) in *Slugt78d-b* pollen supplemented with 200 μM flavonols or an equivalent amount of DMSO (mock). **(E-H)** Each data point is represented by a circle on the violin or bar graphs. Statistical significance was determined using one-way ANOVA followed by Dunnett’s multiple comparisons test, with adjusted P-values displayed above the comparisons.

Consistently, we detected *SlF3’H* expression in our leaf sample but not in pollen (Fig. 2F). This suggests that distinctive pollen-specific flavonol profile is shaped through the combined transcriptional regulation of key biosynthesis enzymes. Specifically, the repression of *F3’H* leads to the predominance of kaempferol, while the activated *SlUGT78D-B* and the repression of *SlUGT78D-A* result in enriched kaempferol galactosides in pollen (Fig. 5C and 2F).

To determine whether these pollen-specific flavonols play any roles in pollen tube development, we conducted chemical complementation tests by supplying flavonols to germinating *Slugt78d-b* pollen. When K-gal and Q-gal were supplied to the *Slugt78d-b* pollen, K2 and Q2 were detected, whereas the mock control did not produce K2 and Q2 (Fig. 5d), suggesting that both K-gal and Q-gal were successfully converted to K2 and Q2, respectively, in pollen. Notably, K-gal supplementation increased pollen tube length and reduced the number of ruptured pollen tubes in *Slugt78d-b* (Fig. 5, E and H). However, the same concentration of Q-gal did not alleviate the defective pollen phenotypes (Fig. 5, E and H). We also tested various concentrations of their aglycones, kaempferol and quercetin, but none of them improved the defective pollen phenotypes (Fig. 5, F to H), which further confirms that kaempferol aglycone is not a limiting factor for pollen tube growth, as its levels in *Slugt78d-b* pollen were seven times higher than in the wild type (Fig. S10B).

Taken together, our findings that not all flavonols, but a specific type of flavonol glycoside, effectively rescue defective pollen tube growth in tomato flavonol mutants suggest physiological significance of individual flavonol structure. This may also provide a functional explanation for the evolutionary loss of quercetin galactosides in certain Solanaceae species.

## Discussion

Our study reveals organ-specific flavonol profiles in tomato and highlights the necessity of pollen-specific kaempferol galactosides for proper pollen tube growth, which cannot be compensated by other aglycones or quercetin galactosides. Multiple studies have reported abnormal plant growth and development in flavonol deficient mutants in various species, often attributed to impaired ROS scavenging capacity or to altered auxin transport(*33*, *38*, *60*). Our data demonstrating the indispensable roles of individual flavonol structures suggests an additional layer of physiological significance beyond their collective general antioxidant properties. This organ-specific flavonol accumulation pattern in tomato is achieved through the tight transcriptional regulation of key flavonol biosynthesis enzymes, combined with the constrained sugar specificity of glycosyltransferases. Millions of specialized metabolites have been identified in plants and glycosylation is a common modification across various phytochemicals(*65–67*). Beyond enhancing metabolite stability or solubility, glycosylation may confer specialized functions, potentially evolving to optimize plant fitness in response to environmental changes.

In tomato, 3-*O*-glucosylated flavonols are broadly distributed across various organs and tissues, but kaempferol galactosides, specifically kaempferol 3-*O*-glucosyl(1 → 2)galactoside (K2), are almost exclusively enriched in pollen and not detected in most organs like leaves and fruits (Fig. 1, A and B). This unique flavonol profile correlates with the distinct expression patterns of flavonol 3-*O*-glycosyltransferases (UGTs) and F3’H, the flavonol 3’-hydroxylase necessary for quercetin production. SlUGT78D-B having strict sugar specificity toward galactose is expressed almost exclusively in pollen, whereas SlUGT78D-A having both flavonol 3-*O*-glucosyltransferase and galactosyltransferase activities and F3’H are ubiquitously expressed across all tissues we tested except for pollen grains (Fig. 2, B and C). This differential distribution of flavonol metabolites and their corresponding biosynthesis enzymes implies finely tuned regulations of these metabolic pathways and possible specialized functions of each individual compound in specific organs, tissues, or cells. Several transcription factors have been reported to control the expression of core flavonoid biosynthetic genes, such as chalcone synthase (CHS) and flavonol synthase (FLS) in Arabidopsis(*68–70*). The identity and action modes of transcription regulators responsible for the organ-specific expression of flavonol modification enzymes F3’H and F3GTs remain unknown. But, tight regulation of flavonol modification across organs further implies significance of each flavonol structure.

Although our phylogenetic study did not identify any other UGTs in the F3GT clade besides SlUGT78D-A and B (Fig. 2A), *Slugt78d-a* leaves still accumulated about 20% of glucosylated flavonols despite the loss of functional SlUGT78D-A, indicating that other UGTs may act as flavonol 3-*O*-glucosyltransferases. A previous study reported that 29 out of 91 tested Arabidopsis UGTs showed quercetin glucosyltransferase activities(*48*). Given that SlUGT78D-B lacks 3-*O*-glucosyltransferase activity and SlUGT78D-A is not expressed in pollen, flavanol glucosides (K4 and Q4) detected in pollen must be imported from other organs like tapetum(*71*) or other UGTs having F3Glu activity must exist in pollen. Interestingly, K2 was undetectable in the pollen of *Slugt78d-b* under our growth conditions. Apparently, SlUGT78D-B is the only kaempferol 3-*O*-galactosyltransferase in pollen and no other promiscuous UGTs can compensate for its activity in pollen.

Our amino acid swapping experiment identified key amino acid residues of SlUGT78D-B restricting its sugar specificity. Among the four amino acids (Ala14, Leu19, Ser138, and His373) predicted to face to the sugar binding pocket (Fig. 3), Leu19, Ser138, and His373 are critical to restrict F3Glu activity as modifying one of them enabled SlUGT78D-B to gain F3Glu activity. Despite the relaxed reversibility of sugar specificity, SlUGT78D-B remains constrained from acquiring F3Glu activity while retaining its galactosyltransferase activity, reflecting an adaptive advantage conferred by galactosylated kaempferol in pollen development.

While some Solanaceae species accumulate both K2 and quercetin 3-*O*-glucosyl(1 → 2) galactoside (Q2) in their pollen, Q2 is absent in chili pepper, potato, and tomato (Fig. 5A). The loss of Q2 occurred approximately 23.9 million years ago, coinciding with the separation of the clade of chili pepper, potato, and tomato (Fig. 5B). Given that Q2 did not rescue the defective pollen tube growth of the *Slugt78d-b* mutant (Fig. 5, E and H), Q2 may have been selectively phased out.

K2 is conserved in Solanaceae, not all plants produce K2 in their pollen. However, each species accumulates unique flavonols in their pollen. For example, Arabidopsis and Cannabis accumulate kaempferol 3-*O*-glucosyl(1 → 2)glucoside (kaempferol sophoroside) or quercetin sophoroside in their pollen. Similarly, flavonol sophorosides were detected in maize pollen and the pollen of trees belonging to the families Juglandaceae, Betulaceae, Fagaceae, and Oleaceae(*42*, *72*, *73*) (*10*, *72–75*). Although these pollen-specific flavonol glycosides vary in their decorating sugars (glucoside, galactoside) and aglycones, they have a common 1 → 2 inter-glycosidic linkage (β-1→2) at the 3-OH position of aglycones. It is possible that specific structure(s) of flavonols may have shared function in pollen development.

In general, flavonols are known as antioxidants and reactive oxygen species (ROS) scavengers, critical for maintaining ROS homeostasis, which in turn affects pollen development(*59*, *60*, *76*, *77*). Consistently, tomato *f3h* mutant that is deficient in all flavonols showed deformed pollen grains and defective pollen germination and tube growth accompanied by increased ROS levels(*59*, *60*). Supplementation with flavonol aglycones reduced ROS levels and alleviate pollen defects(*59*, *60*), suggesting the importance of flavonols in maintaining pollen integrity. In *Slugt78d-a/b* pollen, the level of kaempferol was seven times higher than in wild type (Fig. S10), yet the pollen showed defective pollen tube growth, which was not restored with further supplementation of various concentration of kaempferol (Fig. 5, F to H). Instead, the specific galactosylated kaempferol is critical for proper pollen tube growth in tomato (Fig. 5, F to H), suggesting the physiological specificity of each flavonol structure-additional layer of flavonol’s function besides their general role as antioxidants. Several studies indicate that flavonols may regulate auxin transport by stabilizing PIN-FORMED (PIN) protein dimers(*26*, *78–81*). It is possible that specific type(s) of flavonols may affect pollen tube growth through directly or indirectly interacting with molecules involving in auxin transport (*2*, *35*, *82*).

Recently, flavonol transporters were identified in rice and Arabidopsis(*54*, *83*). In Arabidopsis, Flavonol Sophoroside Transporter 1 (FST1) mediates localization of flavonol glycosides(*54*). As it expresses in tapetum, flavonol soporosides available in tapetum can be transported to pollen via FST1 and Solyc03g113440 is a putative FST1 homolog in tomato(*54*). Interestingly, *Solyc03g113440* expresses exclusively in pollen grains (Fig. S13), and its role remains unknown. Tomato pollen does not accumulate flavonol sophorosides, but a kaempferol galactoside K2 is abundant in pollen. K2 seems to be synthesized predominantly in pollen grains rather than transported substantially from other organs or tissues, as K2 biosynthesis genes are expressed in pollen grains and K2 is absent in pollen grains of *Slugt78d-b* mutants although SlUGT78D-A having flavonol 3-*O*-galactosyltransferase activity functions ubiquitously in other organs of *Slugt78d-b*. The tomato FST1 homolog may potentially transport flavonol glycosides such as K2 or Q4 in pollen.

In conclusion, the exclusive accumulation of pollen-specific flavonol galactosides, resulted from differentially expressed *F3’H* and *F3GTs* having distinctive sugar specificities, reflects a regulatory network finely tuned to meet the developmental needs of pollen. This study demonstrates that specific flavonol galactosides rather than overall abundance of total flavonols matter for proper pollen tube growth and male fertility in tomato. Moreover, the loss of quercetin galactosides in certain Solanaceae species suggests there are selective pressures to shape flavonol composition in pollen, favoring the functional advantages conferred by kaempferol galactosides. These insights advance our understanding of flavonol function in reproductive development and may inform future strategies aimed at enhancing pollen viability and seed yield through targeted manipulation of pollen specific flavonol glycosylation (Fig. 6).

**Fig. 6.**
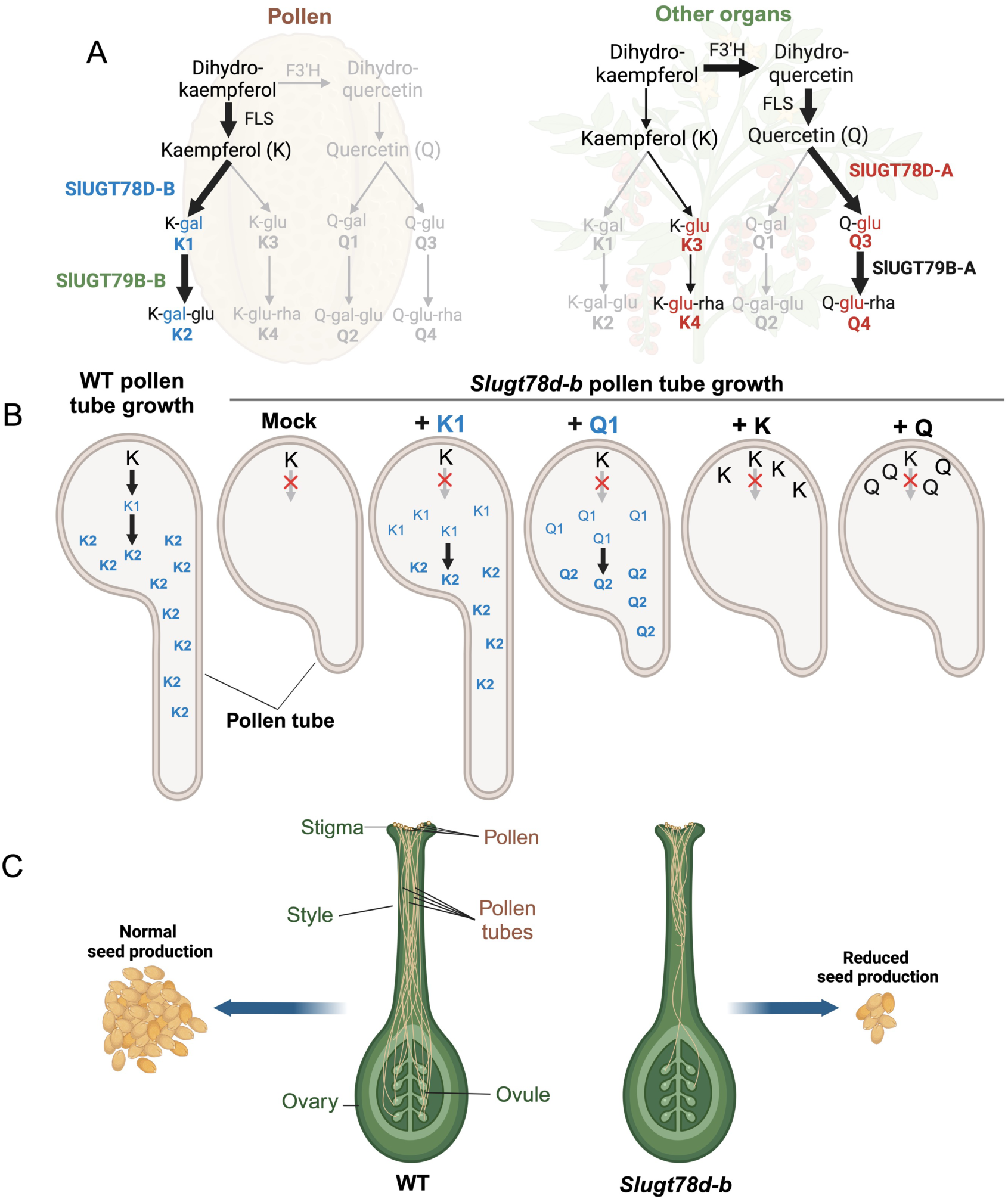
Working model depicting the role of flavonol glycosylation in successful fertilization. **(A)** Organ-specific flavonol glycoside biosynthesis pathways in tomato pollen and other organs. In pollen, kaempferol (K) is synthesized from dihydrokaempferol via flavonol synthase (FLS). The glycosyltransferase SlUGT78D-B catalyzes the formation of kaempferol 3-*O*-galactoside (K-gal, K1), which is further modified by SlUGT79B-B to produce kaempferol 3-*O*-galactoside-2’’-*O*-glucoside (K-gal-glu, K2), a pollen-specific flavonol glycoside. In other organs, kaempferol (K) and quercetin (Q) are produced via FLS, with F3’H catalyzing the conversion of dihydrokaempferol to dihydroquercetin, a precursor of quercetin. These flavonols are glycosylated by SlUGT78D-A to form kaempferol 3-*O*-glucoside (K-glu, K3) and quercetin 3-*O*-glucoside (Q-glu, Q3). Further modification by SlUGT79B-A produce kaempferol 3-*O*-glucosyl(1→6)rhamnoside (K-glu-rha, K4) and quercetin 3-*O*-glucosyl(1→6)rhamnoside (Q-glu-rha, Q4), which are abundant in vegetative tissues. **(B)** Schematic illustration of pollen tube growth defects in *Slugt78d-b* mutants and chemical complementation analysis. Wild-type (WT) pollen accumulates kaempferol (K) and its glycosylated derivatives, including K1 (kaempferol 3-*O*-galactoside) and K2 (kaempferol 3-*O*-galactoside-2’’-*O*-glucoside), which are essential for normal pollen tube growth. In *Slugt78d-b* mutant pollen, the absence of K1 and K2 results in defective pollen tube growth, as observed in the mock treatment. Chemical complementation was conducted by supplementing *Slugt78d-b* mutant pollen with various flavonol derivatives. Among the tested compounds, only the addition of K1 successfully rescued the pollen tube growth defect by restoring K2 biosynthesis. In contrast, complementation with Q1 (quercetin 3-*O*-galactoside), K, or Q failed to restore normal pollen tube growth. **(C)** Schematic illustration of reduced seed production caused by defective pollen tube growth in *Slugt78d-b* mutants. In WT plants, pollen tubes successfully germinate and grow through the style, delivering male genetic material to the ovules in the ovary, resulting in normal seed production. In contrast, *Slugt78d-b* mutant pollen exhibits defective pollen tube growth, impairing fertilization and significantly reducing seed production. This schematic highlights the critical role of *SlUGT78D-B* in regulating pollen tube development and ensuring efficient seed production in tomato.

## Materials and Methods

### Plant material and growth conditions

The Micro-Tom variety was used as the wild-type plant and for generating CRISPR-Cas9-mediated mutant lines. All tomato plants were germinated in 3.7 x 3.7 x 5.8 cm seedling trays and then transplanted into 9 x 9 x 9 cm individual pots 3 weeks after germination, where they were grown until fruit and seed production. Tomato plants were grown at 22°C ± 1°C under a 16-hour day/8-hour night photoperiod.

### Generation of CRISPR-Cas9-mediated mutant lines

To disrupt SlUGT78D-A and SlUGT78D-B using the CRISPR-Cas9 system, gRNAs targeting both genes were designed using the CHOPCHOP software (https://chopchop.cbu.uib.no/)(84). Specifically, two 20 bp gRNAs (5′-ATTTGGGATGGCGTCAAACA-3′) and (5′-CCAAAGGGTCACGTGCTCGG-3′) were selected for *SlUGT78D-A* and *SlUGT78D-B,* respectively. Then, a DNA fragment containing a BsaI restriction site, the *SlUGT78D-A* gRNA, a gRNA scaffold, the U6-26t and U6-6P sequences, the *SlUGT78D-B* gRNA, and another BsaI restriction site was synthesized (Twist Biosciences, South San Francisco, CA, USA), which was cloned into the binary vector pHSE401 (Addgene, #62201) using the BsaI restriction site(*85*). The recombinant pHSE401 vector was then transformed into *A. tumefaciens* strain GV3101 and inoculated into cut cotyledons following a tomato transformation method^39^. The mutant lines were selected according to the sequencing results of PCR products amplified with primers no. 1 and 2 for *SlUGT78D-A*, and no. 3 and 4 for *SlUGT78D-B* (Table S3).

### Phylogenetic analysis

Putative tomato UGTs were identified by querying the UniProt database using “UDP-glycosyltransferase” as the search keyword, with *Solanum lycopersicum* (taxon ID: 4081) specified as the organism. Candidate sequences were further screened for the presence of the PSPG (Plant Secondary Product Glycosyltransferase) domain using the NCBI Conserved Domains database (https://www.ncbi.nlm.nih.gov/Structure/cdd/wrpsb.cgi). Sequences lacking the PSPG domain were excluded, resulting in a final set of 182 amino acid sequences for phylogenetic analysis. Previously characterized F3GTs and F3GGTs from other species (Table S4) were also included in our phylogenetic analysis. All sequences were aligned using the MUSCLE algorithm. Phylogenetic analysis was performed using the maximum likelihood method with the Goldman +Freq. model and Gamma distribution with 5 categories in MEGA11 software, with 1000 bootstrap replicates. Solyc12g088170 (anthocyanin acyltransferase; SlFdAT1)(*87*) was used for the outgroup in the phylogenetic analysis.

### Metabolite analysis using HPLC and LC-MS

Plant samples, including pollen samples from other Solanaceae plants, were collected and immediately frozen in liquid nitrogen. Soluble metabolites in the frozen samples were extracted using 50% methanol at a concentration of 50 mg fresh weight/ml, except for pollen, which was prepared to a concentration of 28.5 mg fresh weight/ml after 15 minutes of sonication at room temperature. The extracts were centrifuged at 10,000 g for 10 minutes to remove plant tissue and insoluble debris before being used for metabolite analysis. Authentic standards used for metabolite identification are kaempferol, kaempferol 3-*O*-glucoside, kaempferol-3-*O*-galactoside, kaempferol 3-*O*-rhamnosyl(1→6)glucoside, quercetin 3-*O*-glucoside, and quercetin 3-*O*-galactoside, which were obtained from Chengdu Biopurify Phytochemicals Ltd. (Chengdu, China); quercetin and quercetin 3-*O*-rhamnosyl(1→6)glucoside from Sigma-Aldrich (St. Louis, MO, USA); and chlorogenic acid from TCI (Portland, OR, USA).

For HPLC analysis, the extracts were analyzed using an UltiMate 3000 HPLC system (Thermo Fisher Scientific, Waltham, MA, USA) equipped with an Acclaim™ RSLC120 C18 column (100 mm x 3 mm, 2.2 µm) (Thermo Fisher Scientific, Waltham, MA, USA). The mobile phase consisted of solvent A (0.1% formic acid in water) and solvent B (100% acetonitrile) with the following linear gradient: 5-14% B for 2.2 minutes, 14-18% B for 9 minutes, and 18-95% B for 3.5 minutes, with a flow rate of 0.5 ml/min. Flavonoid contents were quantified based on peak area at 345 nm using authentic standards. K-3-*O*-gal-2’’-*O*-glu (K2) was quantified equivalently to K-3-*O*-glu-6’’-*O*-rha.

For Semi-targeted LC-MS analysis, an Ultimate 3000 UHPLC coupled to a TSQ Quantiva triple quadrupole mass spectrometer was used (Thermo Fisher Scientific, Waltham, MA, USA). 3,4,5-trihydroxycinnamic acid from Sigma-Aldrich (St. Louis, MO, USA) was used as an internal standard. The injection volume for all samples was 5 μl. Analytes were eluted on an Agilent InfinityLab Poroshell HPH-C18 column (2.1 × 150 mm, 1.9 µm particle size) at a column temperature of 40°C using 0.1% formic acid in water (mobile phase A) and 0.1% formic acid in acetonitrile (mobile phase B). A gradient elution at a flow rate of 0.4 ml/min was applied as follows: 2–42% B for 28 min, 42–100% B for 2 min. The mass spectrometer operated in negative ionization mode. The ESI parameters were as follows: spray voltage (positive), 3500 V; spray voltage (negative), 2500 V; ion transfer tube temperature, 325°C; vaporizer temperature, 275°C; sheath gas, 35 Arb; aux gas, 10 Arb; and sweep gas, 0 Arb. Selective reaction monitoring (SRM) was used for metabolite detection. MS/MS parameters were optimized for each metabolite using flow injection analysis of individual standards. Collision-induced dissociation (CID) gas pressure was set at 2 mTorr with a dwell time of 0 ms, and quadrupole resolution (FWHM) was kept at 0.2. Data processing was performed using Xcalibur v4.0 (Thermo Fisher Scientific, Waltham, MA, USA).

To identify the unknown flavonol glycoside (Peak1; K2) lacking an authentic standard shown in Figure 1A, a combination of HPLC and LCMS was employed. K2 was isolated based on its retention time on an HPLC system, and the isolated peak was used for further analysis.

Specifically, a Thermo Scientific™ Q Exactive Focus mass spectrometer coupled with a Dionex™ Ultimate™ RSLC 3000 UHPLC system was used for the analysis. The system was equipped with an H-ESI II probe on an Ion Max API Source. Analytes were separated on an Agilent Poroshell 120 EC-C18 column (2.7 μm, 3.0 × 50 mm) using solvent A (water with 0.1% formic acid) and solvent B (acetonitrile with 0.1% formic acid). The LC program with a flow rate of 0.5 ml/min was as follows: 10% B for 2 min, 10-95% B over 8.5 min, 95% B for 2.5 min, 95% to 10% B in 0.5 min, followed by re-equilibration at 2% B for 2 min. MS signals were acquired under Full MS positive ion mode, covering a mass range of m/z 150-2000, with a resolution of 35,000 and an AGC target of 1E6. The precursor ion (m/z 611.1597) was selected in the Orbitrap, typically with an isolation width of 3.0 m/z and fragmented in the HCD (Higher-energy C-trap dissociation) cell with stepwise a collision energy (CE) of 15.

### Ubiquinone quantification

Ubiquinone extraction and quantification in tomato samples were modified from Ducluzeau et al(*88*). Tomato pollen samples, 0.8-1.8 mg of pollen grains, were collected and spiked with 3 nmoles of ubiquinone-9. The samples were homogenized in 0.4 ml of 95% (v/v) ethanol using a Pyrex tissue grinder. The grinder was rinsed twice with 0.3 ml of 95% (v/v) ethanol, and washes were combined to the original extract in a 15-ml Pyrex tube containing 0.5 ml of water. The mixture was partitioned twice with 5 ml hexane. Hexane layers were combined, evaporated to dryness with gaseous nitrogen, and resuspended in 0.2 ml of ethanol:dichloromethane (10:1, v/v). Due to the pigment complexity of the pollen extracts, it was not possible to fully resolve the ubiquinol-10 peak. Quinol forms were therefore converted to their respective quinone counterparts by adding 1 μl of 15% H_2_O_2_. Samples were then briefly centrifuged (5 min; 18,000 g) and immediately analyzed by HPLC on a 5 μm Supelco Discovery C-18 column (250 × 4.6 mm, Sigma-Aldrich) thermostated at 30°C and developed in isocratic mode at a flow rate of 1 ml/min with methanol:hexane (90:10, v/v). Ubiquinone forms were detected spectrophotometrically at 275 nm. Retention times were 13.5 min for ubiquinone-9 and 18 min for ubiquinone-10 (Q10). Data were corrected for recovery of the ubiquinone-9 internal standard and recovery values ranged from 79% to 95%.

### RNA extraction and expression analysis

Tomato samples were collected and immediately frozen in liquid nitrogen. The samples were then homogenized using a Benchmark BeadBlaster 24 homogenizer (Benchmark Scientific, Edison, NJ, USA). Total RNA was extracted using TRIzol™ reagent (Life Technologies Inc., Gaithersburg, MD, USA).

cDNA synthesis was performed using 2 µg of total RNA and a reverse transcription kit (Thermo Fisher Scientific, Waltham, MA, USA). Quantitative real-time PCR (qRT-PCR) reactions were carried out using a PCR kit (Thermo Fisher Scientific, Waltham, MA, USA). Primer no. 5 and 6 were used for SlUGT78D-A, while primer no. 7 and 8 were used for SlUGT78D-B (Table S3). The expression levels of the target genes were normalized to those of *SlACTIN2* (*Solyc11g005330*) or *SlPP2Acs* (*Solyc05g006590*) as internal controls(*89*, *90*), which were amplified with primer No. 9 and 10 and primer No. 11 and 12, respectively.

### Protein structure prediction

To predict the protein structure of SlUGT78D-B, we employed ColabFold, a computational tool that combines fast homology search with MMseqs2 and the AlphaFold2 algorithm(*91*). The default multiple sequence alignment (MSA) pipeline was used, which included an MMseqs2 search of both the UniRef and environmental sample sequence databases. The resulting predicted protein structure was saved in the Protein Data Bank (PDB) format and visualized using PyMOL 2.5 (Schrödinger, LLC). To determine the ligand position, we overlaid the VvGT1 structure (PDB DOI: https://doi.org/10.2210/pdb2C1Z/pdb), previously resolved via crystallography (*6*), onto the predicted SlUGT78D-B enzyme structure using PyMOL 2.5.

### In vitro enzyme assay

The coding sequences (CDS) of wild-type and mutant *SlUGT78D* genes were synthesized by Twist Biosciences (South San Francisco, CA, USA) and introduced to the pET-28a(+) vector using the NEBuilder HiFi DNA Cloning Kit (NEB, USA). The recombinants were subsequently transformed into *E. coli* strain BL21 using the heat shock method. For the enzyme assay, the *E. coli* harboring the expression vector was cultured in 50 ml of Luria broth (LB) medium at 37°C with shaking at 250 rpm until the optical density at 600 nm (O.D.600) reached approximately 0.5. Isopropyl β-D-1-thiogalactopyranoside (IPTG) was then added to a final concentration of 0.1 mM to induce protein expression. After overnight incubation at 18°C, the cells were harvested by centrifugation at 8000 rpm for 10 minutes at 4°C. The bacterial pellet was resuspended with protein extraction buffer containing 50 mM NaH2PO4·H2O, 300 mM NaCl, 10 mM imidazole, and 0.1% Triton X-100, with the pH adjusted to 8. The resuspended pellet was homogenized using a Benchmark BeadBlaster 24 homogenizer (Benchmark Scientific, Edison, NJ, USA) and centrifuged at 15,000 rpm for 10 minutes to obtain the supernatant. For the enzyme reaction, 100 μl of the crude protein extract was mixed with 100 μl of reaction buffer containing 200 mM Tris-HCl (pH 7.5), 28 mM 2-mercaptoethanol, 200 μM sugar acceptor (flavonol aglycone), and 2 mM UDP-sugar. The total 200 μl reaction mixture was incubated for 30 minutes at 30°C, after which the reaction was stopped by adding 100% methanol. The mixture was then centrifuged for 20 minutes, and 10 μl of the supernatant was injected into HPLC for analysis using the same HPLC running method described in the previous section.

### Pollen tube growth assay

The pollen germination and pollen tube growth assay were conducted following a previously described protocol with minor modifications(*59*, *92*, *93*). Tomato pollen grains were collected from newly opened flowers in the morning and immediately used the same day to preserve viability. The collected pollen was incubated in Pollen Germination Medium (PGM) consisting of 24% (w/v) PEG4000, 0.01% (w/v) boric acid, 2% (w/v) sucrose, 20 mM Hepes buffer (pH 6.0), 30 mM Ca(NO_3_)_2_, 1.5 mM MgSO_4_, and 1 mM KNO_3_, on slide glasses topped with cover slips. After 4 hours of incubation at 22°C in the dark, the images of pollen tubes were captured using an Olympus IX81-DSU Spinning Disk Confocal Microscope (Olympus America Inc., Center Valley, PA, USA). To minimize potential discrepancies in incubation times, at least three replicate slides for each condition and genotype was prepared and the order of capturing confocal microscope images was rotated in each replicate and each batch. At least three independent batches were examined. Pollen tube length and the number of ruptured or intact pollen tubes were measured using ImageJ (ver. 1.54g for MacOS). For the flavonol feeding assay, a stock solution of flavonols dissolved in DMSO was added to PGM to achieve the desired final concentration, while a DMSO-only solution was used as a mock control.

## Acknowledgments

We thank Dr. Marcio Resende and Dr. Leo Hoffmann Jr. for sharing potato pollen, Dr. Bala Rathinasabapathi and Dr. Jingwei Fu for allowing us to collect pollen from their pepper and *Nicotiana tabacum* plants, and Dr. Thomas A. Colquhoun and Joo Young Kim for allowing us to collect pollen from their petunia plants. We also thank Dr. Tong Geon Lee for helping us propagate tomato mutant seeds and for advising on informatics. We also thank Dr. Ru Dai and Julia Ball for helping with tomato seed collection.

## Supplementary Materials

**Figure S1.**
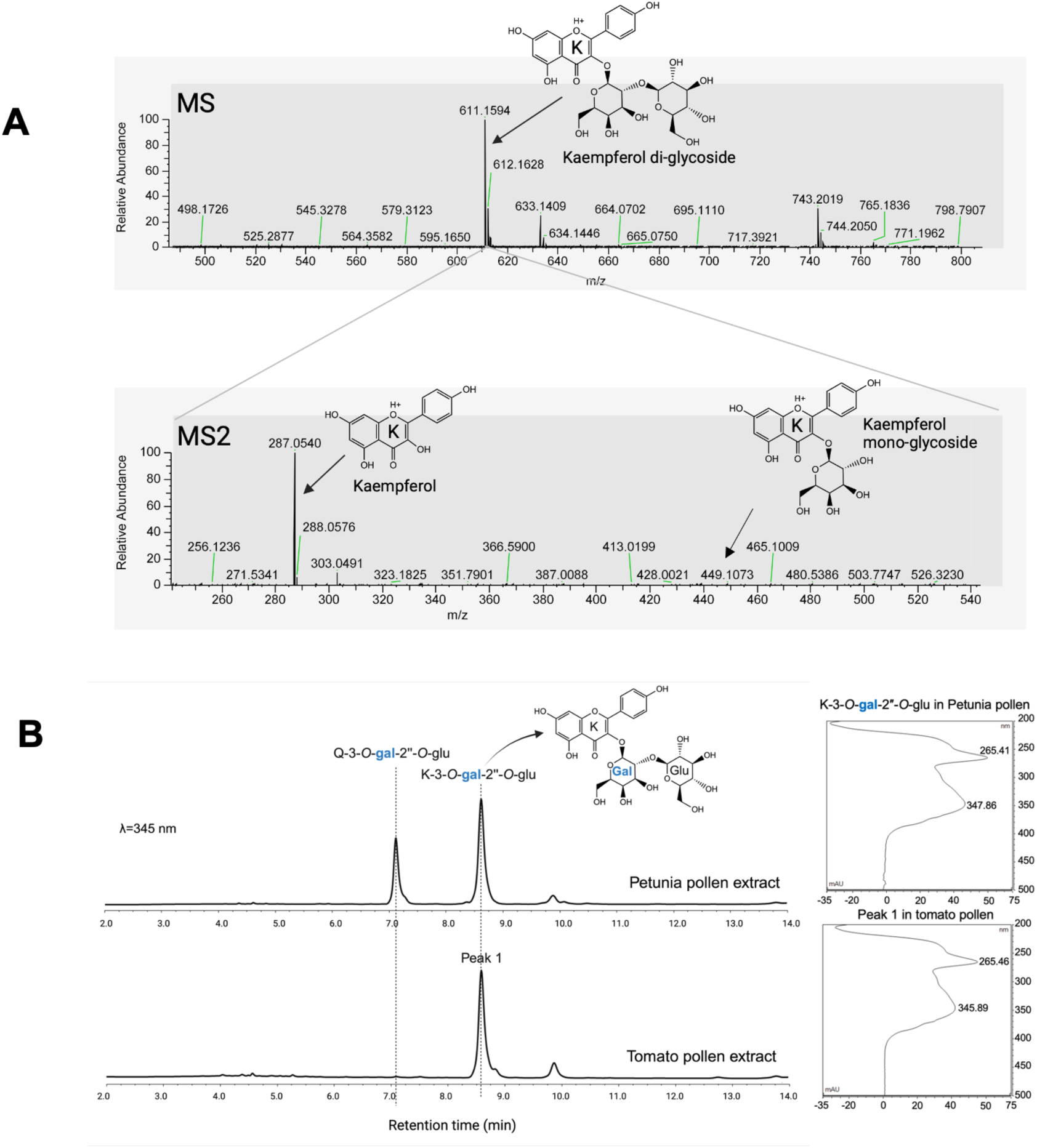
Identification of a pollen specific kaempferol 3-*O*-glucosyl(1 → 2)galactoside in tomato pollen grains. **(A)** LC-MS analysis of the isolated peak from HPLC, which predominantly appears in pollen. MS and MS2 spectra indicate it as a kaempferol di-glycoside containing two hexose units. The MS spectrum displays the precursor ion in positive mode with an m/z of 611.1594, corresponding to kaempferol di-glycoside. The calculated exact mass is 610.1521 Da, determined by subtracting the proton mass (1.0073 Da) from the m/z value. The MS2 spectrum shows the fragmentation pattern, highlighting the major fragment ions, including kaempferol (m/z 287.0540) and kaempferol galactoside or glucoside (m/z 465.1009). **(B)** HPLC chromatograms and UV spectra of pollen extracts from Petunia and tomato. The UV spectra and retention time of the tomato pollen peak (Peak 1) match with the most abundant flavonol of petunia pollen, which was identified as either kaempferol or quercetin 3-*O*-glucosyl(1 → 2)galacoside (K/Q-3-*O*-gal-2’’-*O*-glu) in a previous report(*1*). Taken together with mass data and profile comparison with Petunia pollen, we concluded that Peak 1 is kaempferol 3-*O*-glucosyl(1 → 2) galactoside (K2; K-3-*O*-gal-2’’-*O*-glu).

**Figure S2.**
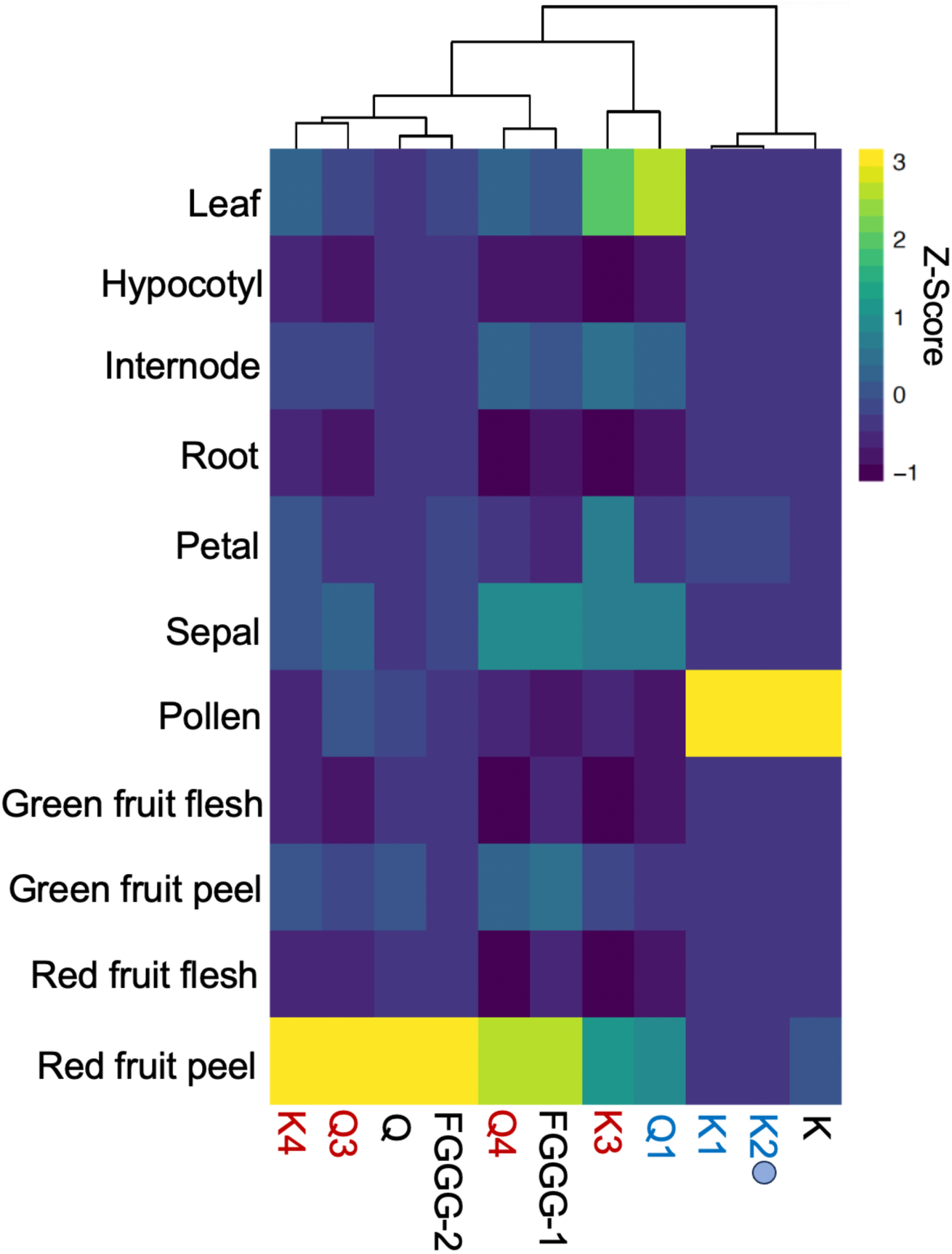
Heatmap of flavonol glycosides across tomato organs detected with LC-MS. The heatmap displays the relative abundance (Z-score) of various flavonol glycosides and two aglycones, K and Q, across different tomato organs. Each column represents a specific flavonol, and the Z-score indicates its relative abundance across different tomato organs. K, K1, and K2 are almost exclusively detected in pollen grains. K2 is the compound with a calculated exact mass of 610.1534, corresponding to the Peak 1, identified in Supplementary Figure S1. The raw data for this LC-MS analysis is available in Supplementary Table S1.

**Figure S3.**
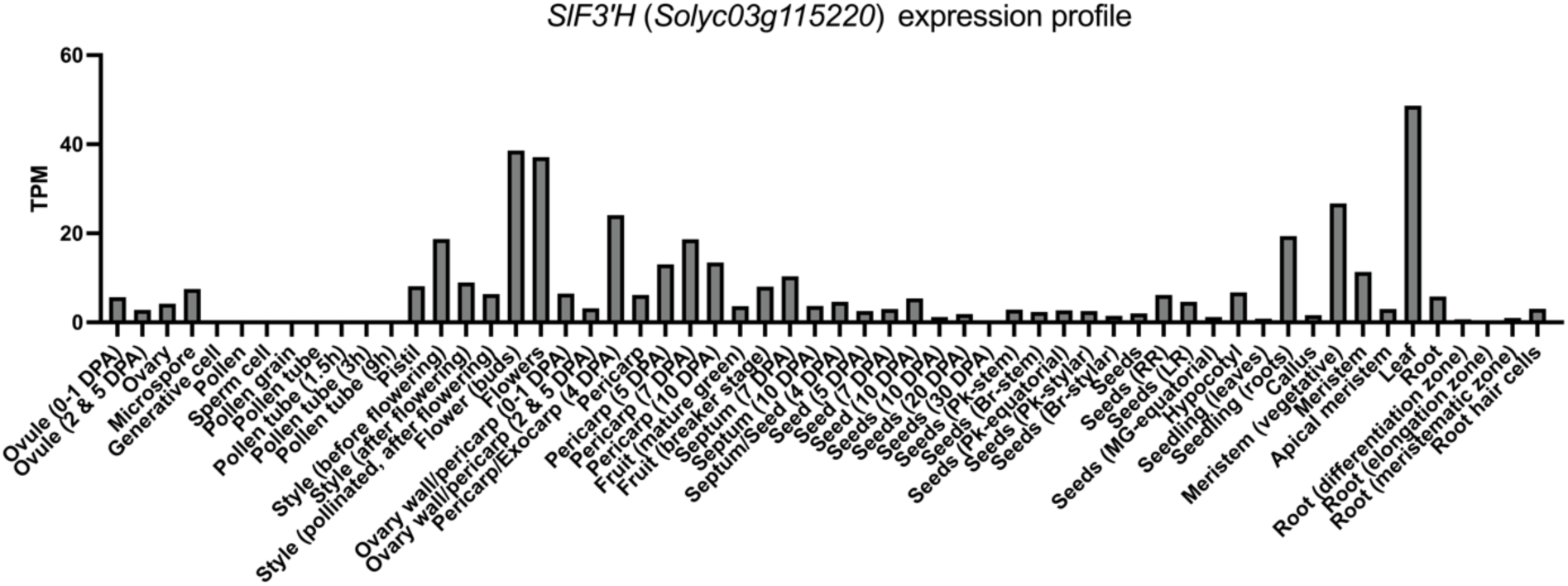
Gene expression profile of *SlF3’H* (*Solyc03g115220*). The raw transcript per million (TPM) data were obtained from the public transcriptome database CoNekT (https://conekt.sbs.ntu.edu.sg). The graph shows the minimal expression of SlF3’H in pollen grains and at different stages of pollen tube development.

**Figure S4.**
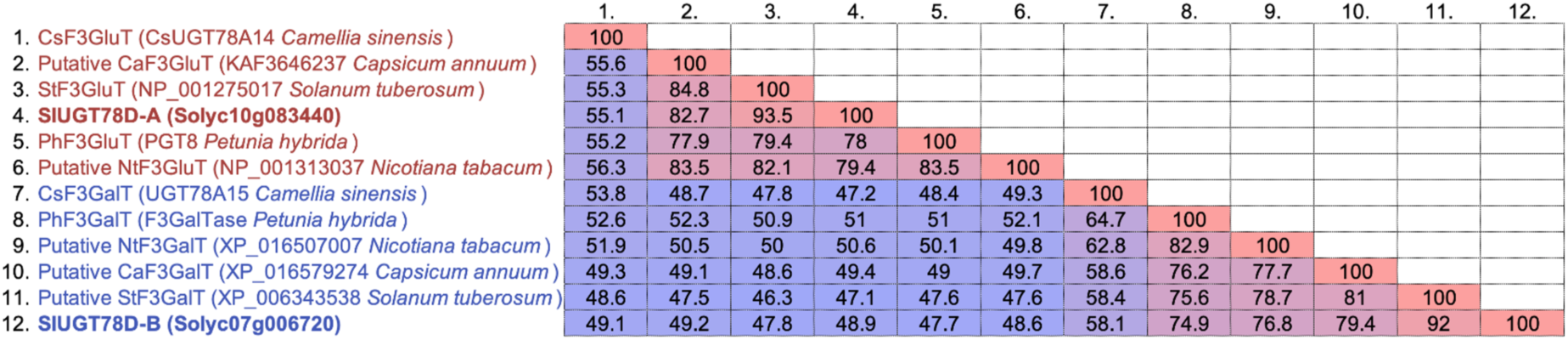
Homologs of flavonol 3-*O*-glycosyltransferase enzymes (SlUGT78D-A and SlUGT78D-B) in other Solanaceae species. A heatmap displaying the percentage identity of protein sequences among SlUGT78D-A, SlUGT78D-B, their homologs in four Solanaceae species, *Nicotiana tabacum*, *Capsicum annuum*, *Solanum tuberosum*, *Petunia x hybrida,* and CsUGT78A14 and CsUGT78A15 from *Camellia sinensis*. Flavonol 3-*O*-glucosyltransferase (CsUGT78A14) and flavonol 3-*O*-galactosyltransferase (CsUGT78A15) from *Camellia sinensis* (tea) are also included in this comparison. High identity percentages suggest functional conservation.

**Figure S5.**
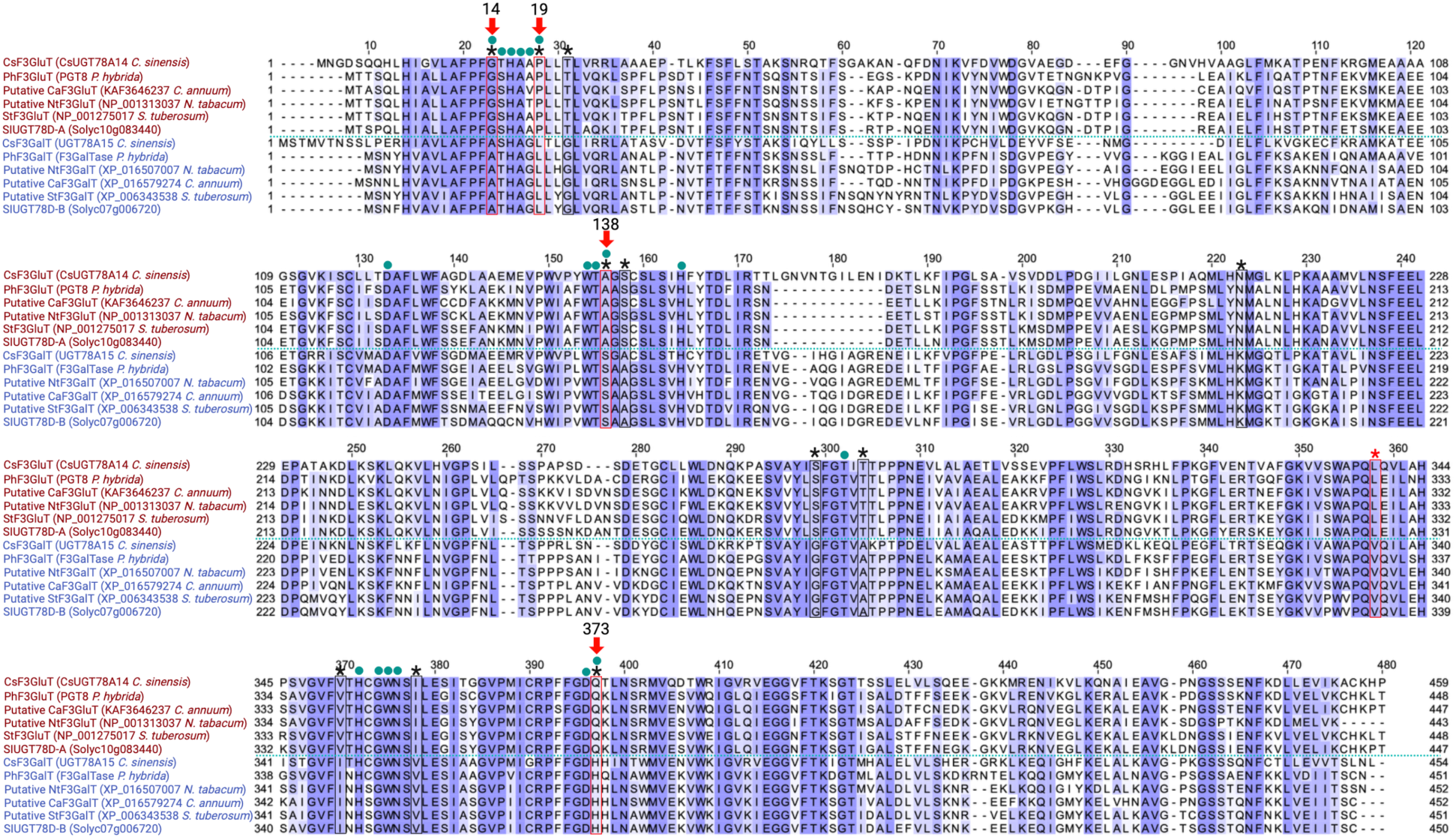
Protein sequence alignment of F3GTs. Sequence alignment of F3GT enzymes from various species, including *Petunia x hybrida, Nicotiana tabacum, Capsicum annuum, Solanum lycopersicum, Solanum tuberosum,* and *Camellia sinensis*. Amino acids highlighted in blue represent conserved residues, based on the BLOSUM62 score. Among these conserved amino acids, 11 amino acids are uniquely conserved in the SlUGT78D-A group and the SlUGT78D-B group, marked with asterisks. Green circles indicate amino acid positions predicted to protrude into ligand binding sites, as inferred from the GT1 structure (*Vitis vinifera*)(*2*, *3*). Red arrows mark the four amino acids identified in this study as essential for sugar specificity.

**Figure S6.**
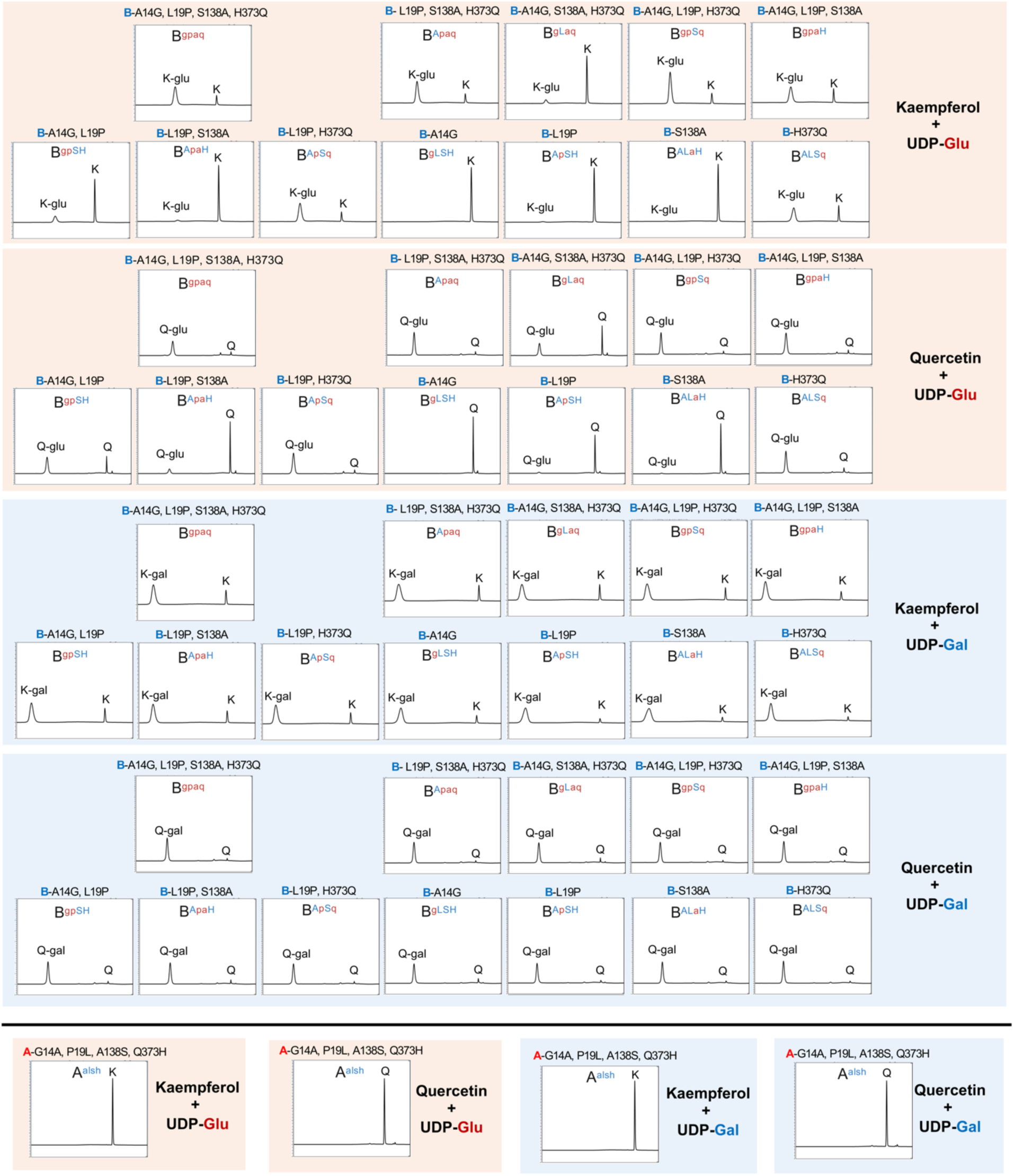
Enzyme activities of modified F3GT enzymes. HPLC chromatograms representing the enzymatic activity of modified F3GT enzymes. The upper panels depict activities of SlUGT78D-B mutant variants, while the lower panels show flavonol 3-*O*-glucosyltransferase or 3-*O*-galactosyltransferase activities of the SlUGT78D-A^alsh^ mutant. The activity was tested using two substrates, kaempferol (K) or quercetin (Q), in the presence of UDP-glucose (UDP-Glu) or UDP-galactose (UDP-Gal) as sugar donors. Each chromatogram represents a different combination of enzymes, aglycones (K or Q), and sugar donors (Glu or Gal). The chromatograms were obtained at a wavelength of 345 nm, showing the resulting flavonol glycosides produced under each condition. K-glu: Kaempferol 3-*O*-glucoside, K-gal: Kaempferol 3-*O*-galactoside, Q-glu: Quercetin 3-*O*-glucoside, Q-gal: Quercetin 3-*O*-galactoside.

**Figure S7.**
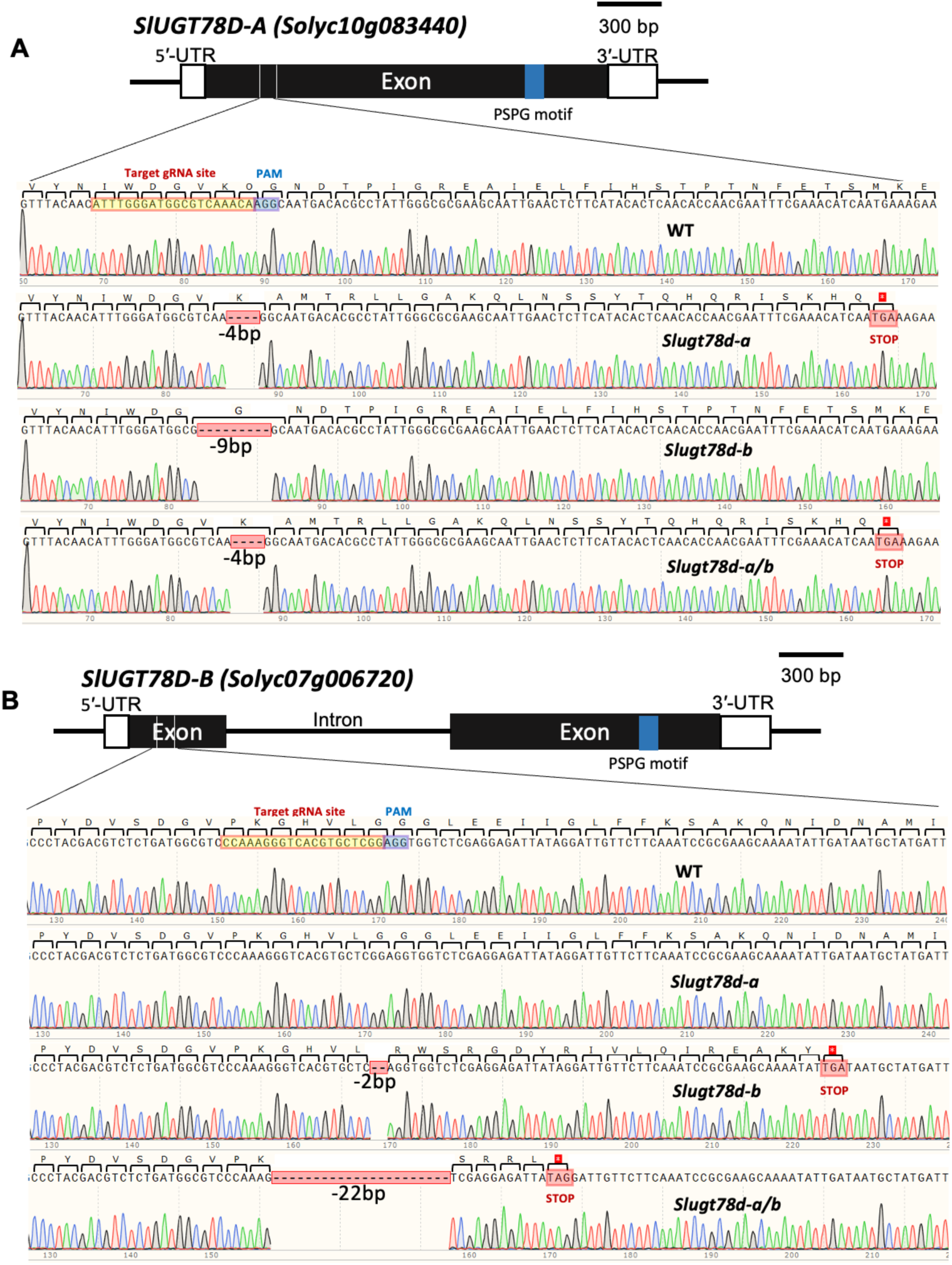
Generation of *Slugt78d-a*, *Slugt78d-b*, and *Slugt78d-a/b* mutants using the CRISPR-Cas9 system. Schematic gene structures of *SlUGT78D-A* (**a**) and *SlUGT78D-B* (**b**) are shown, along with sequencing chromatograms displaying the regions where gene editing occurred. **(A)** *Slugt78d-a* and *Slugt78d-a/b* mutants have a 4 bp deletion in *SlUGT78D-A*, resulting in an early stop codon. *Slugt78d-b* has a 9 bp deletion in *SlUGT78D-A*, but this mutation does not cause a frameshift. **(B)** For *SlUGT78D-B*, the *Slugt78d-b* mutant has a 4 bp deletion, and the *Slugt78d-a/b* mutant has a 22 bp deletion, both of which result in early stop codons, disrupting the gene function.

**Figure S8.**
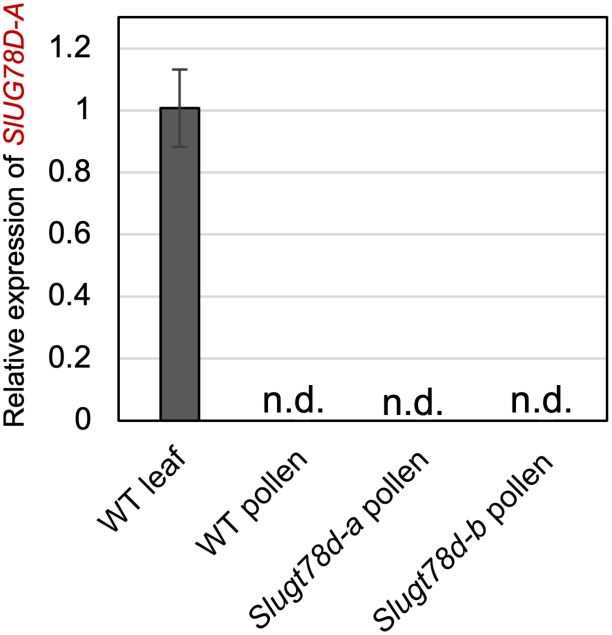
*SlUGT78D-A* is not transcriptionally induced in *Slugt78d-b* pollen. Relative expression of *SlUGT78D-A* in wild-type leaf and pollen grains of WT, *Slugt78d-a,* and *Slugt78d-b*, measured using qRT-PCR. Expression levels were normalized using *SlACTIN2* (*Solyc11g005330*) as an internal control. Data represent mean ± SD (n=4). n.d. indicates not detected.

**Figure S9.**
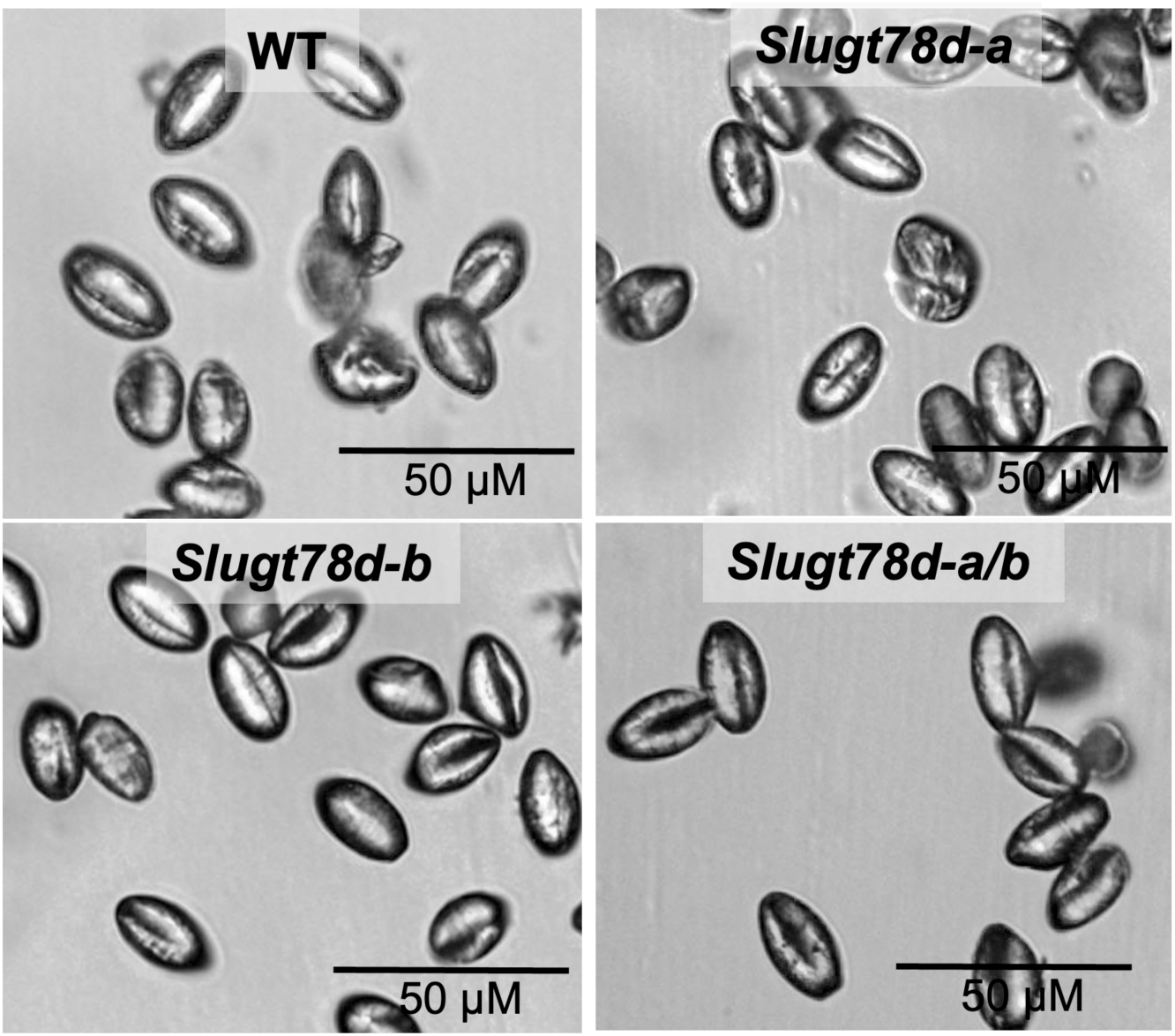
Pollen grain morphology in wild type and three *Slugt78d* mutants. Microscopic images of pollen grains from wild type (WT), *Slugt78d-a*, *Slugt78d-b*, and *Slugt78d-a/b*. The images do not show any obvious morphological alterations in the pollen grains of mutants compared to wild type.

**Figure S10.**
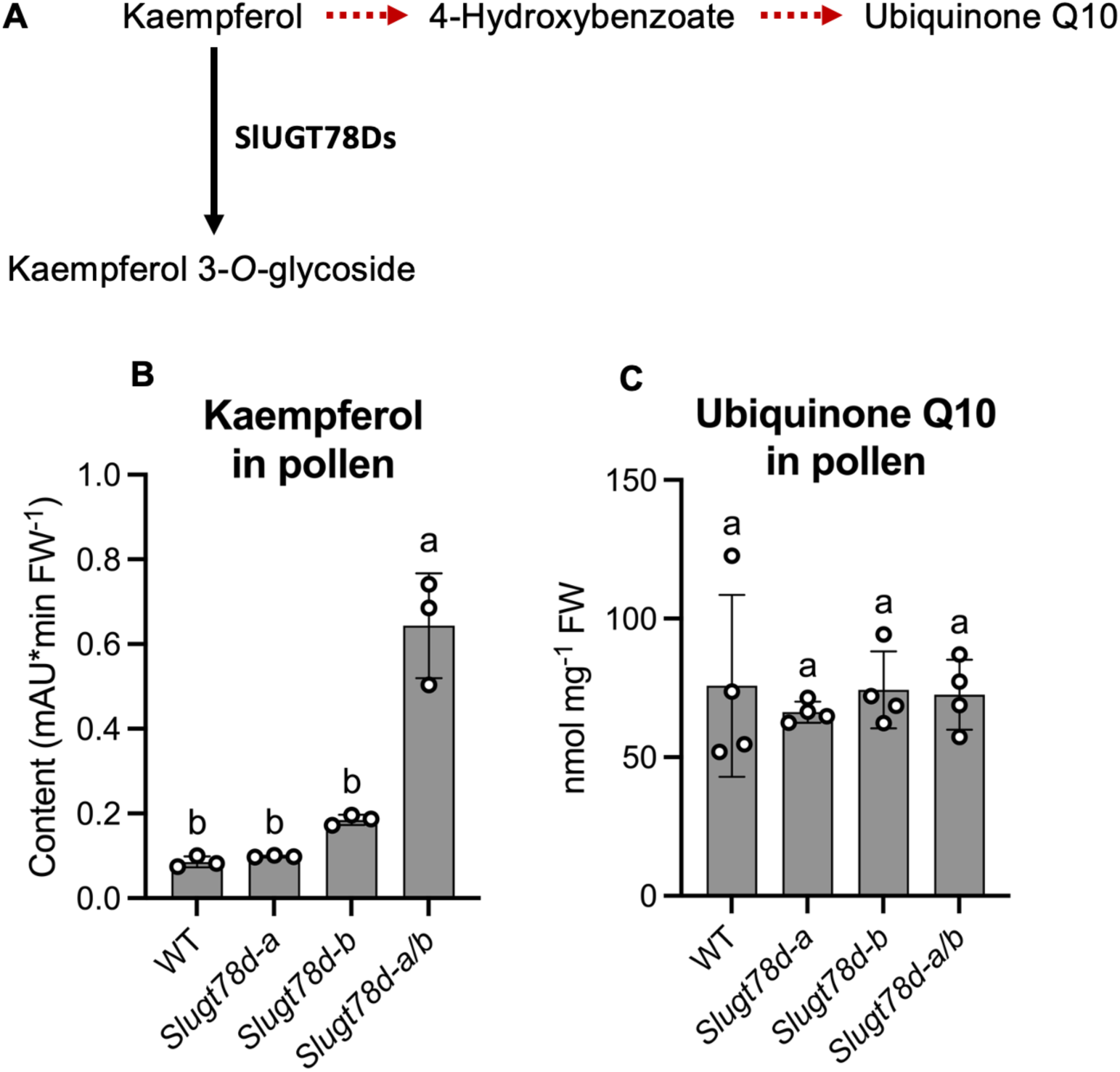
Ubiquinone Q10 content in pollen grains. **(A)** Schematic diagram shows the biosynthetic pathway of ubiquinone Q10 from kaempferol as a precursor. **(B)** The levels of free kaempferol in the pollen grains of WT and three *Slugt78d* mutants. **(C)** The levels of ubiquinone Q10 in the pollen grains of WT and three *Slugt78d* mutants. **(B, C)** Data represent mean ± SD (n=3). Statistical significance was determined using one-way ANOVA, followed by Tukey’s post-hoc test, with different letters indicating significant differences among groups (P-value < 0.05). The metabolite levels were normalized to fresh weight (FW). Each data point is shown as an open circle on the bar graph.

**Figure S11.**
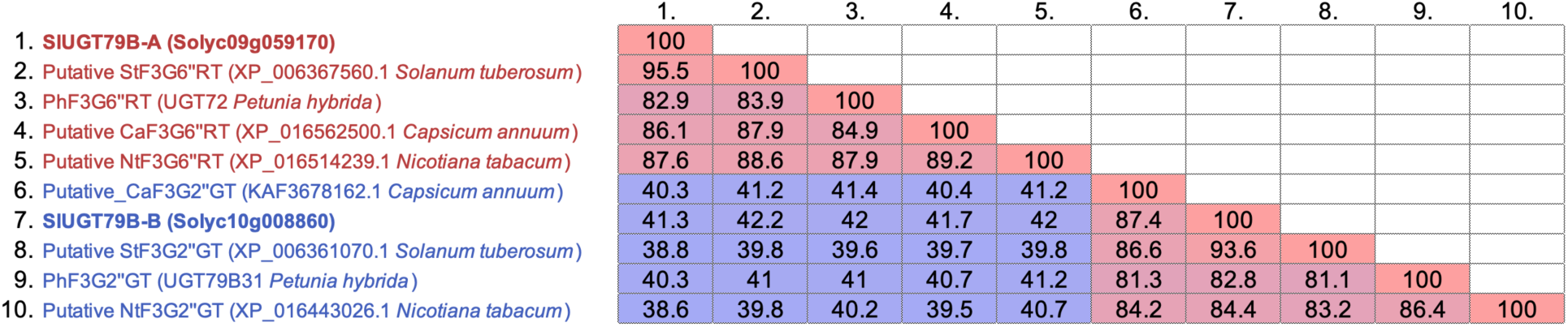
Homologs of flavonol 3-*O*-glycoside: glycosyltransferase (F3GGT) enzymes (SlUGT79B-A and SlUGT79B-B) in other Solanaceae species. A heatmap displaying the percentage identity of protein sequences among SlUGT79B-A, SlUGT79B-B, their homologs in four Solanaceae species, *Nicotiana tabacum*, *Capsicum annuum*, *Solanum tuberosum*, *Petunia x hybrida*. High identity percentages suggest functional conservation.

**Figure S12.**
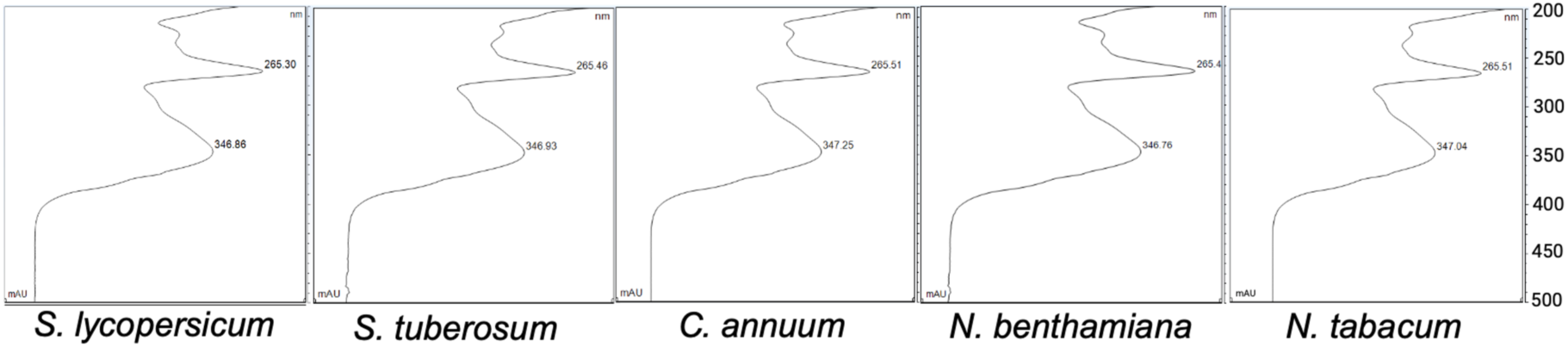
Conservation of pollen-specific kaempferol 3-*O*-glucosyl(1 → 2)galactoside (K2). UV absorbance spectra corresponding to the K2 peaks in pollen extracts from five Solanaceae plants.

**Figure S13.**
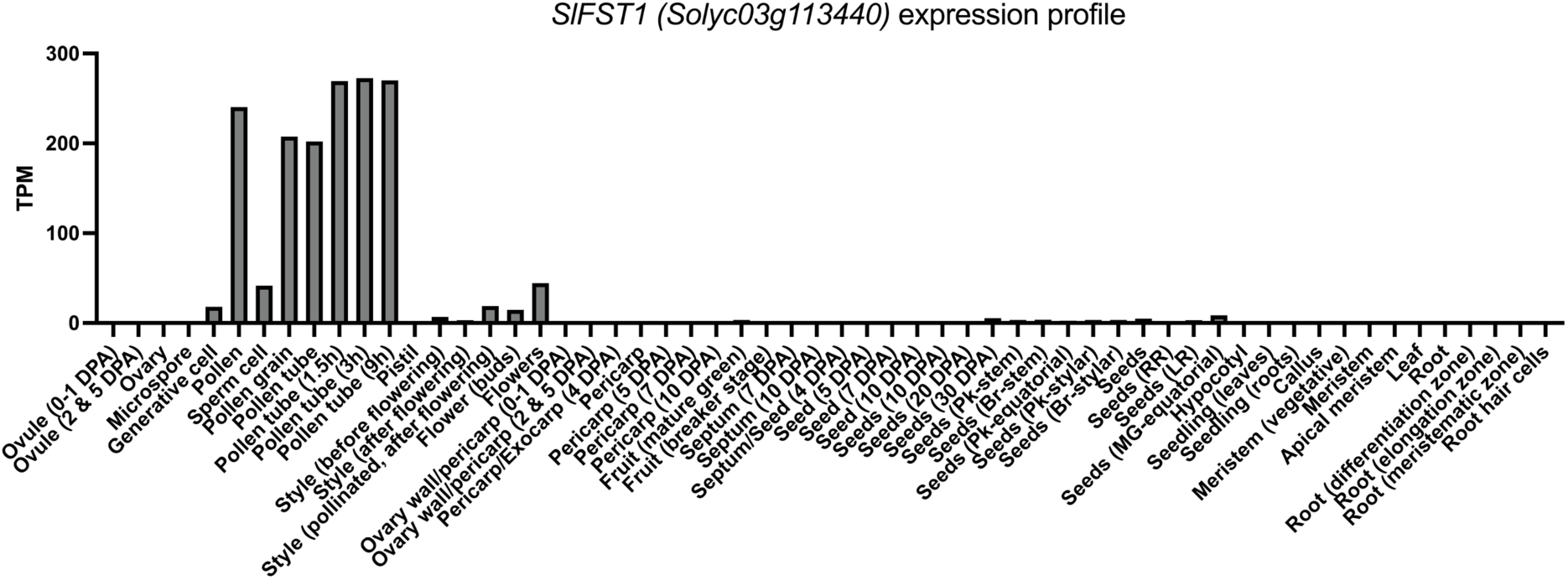
Gene expression profile of *SlFST1* (*Solyc03g113440*), a homolog of Arabidopsis *FST1*. The raw transcript per million (TPM) data were obtained from the public transcriptome database CoNekT (https://conekt.sbs.ntu.edu.sg). The graph shows the specific high expression of *SlFST1* in pollen grains and at different stages of pollen tube development.

**Table S3.**
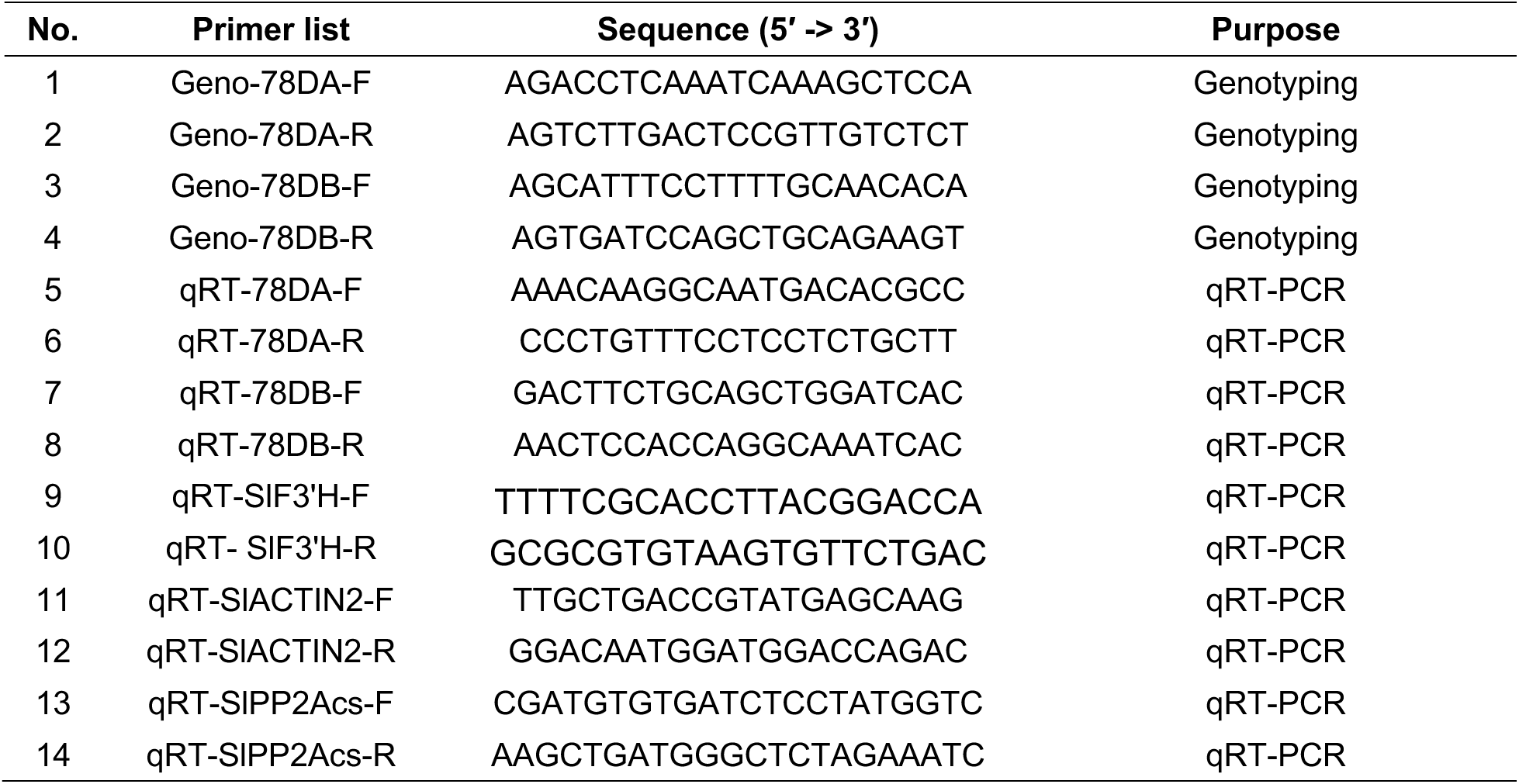
Primer list used in this study.

**Table S4.** Protein accession number for phylogenetic tree.

| Name | Activity | Organism | Accession number |
| --- | --- | --- | --- |
| AtUGT78D1 | F3RhaT | *Arabidopsis thaliana* | AT1G30530.1 (TAIR) |
| AtUGT78D2 | F3GluT | *Arabidopsis thaliana* | AT5G17050.1 (TAIR) |
| AtUGT78D3 | F3AraT | *Arabidopsis thaliana* | AT5G17030.1 (TAIR) |
| VvGT1 | F3GluT | *Vitis vinifera* (Grape) | P51094 (UniProt) |
| VvGT5 | F3GAT | *Vitis vinifera* (Grape) | C7G3B5 (UniProt) |
| VvGT6 | F3Glu/GalT | *Vitis vinifera* (Grape) | C7G3B6 (UniProt) |
| PhF3GalTase | F3GalT | *Petunia x hybrida* | Q9SBQ8 (UniProt) |
| PhPGT8 | F3GluT | *Petunia x hybrida* | Q9SBQ3 (UniProt) |
| Sm3GT | F3GluT | *Solanum melongena* (Eggplant) | Q43641 (UniProt) |
| CsUGT78A14 | F3GluT | *Camellia sinensis* (Tea) | A0A109P632 (UniProt) |
| CsUGT78A15 | F3GalT | *Camellia sinensis* (Tea) | A0A0X9I0C6 (UniProt) |
| ZmBZ1 | F3GluT | *Zea mays* (Corn) | P16165 (UniProt) |
| HvBZ1 | F3GluT | *Hordeum vulgare* (Barley) | P14726 (UniProt) |
| AtUGT79B1 | F3G2"XylT | *Arabidopsis thaliana* | AT5G54060.1 (TAIR) |
| AtUGT79B6 | F3G2"GluT | *Arabidopsis thaliana* | AT5G54010.1 (TAIR) |
| PhUGT79B31 | F3G2"GluT | *Petunia x hybrida* | A0A2Z6FLE1 (UniProt) |
| PhUGT72 | F3G6"RhaT | *Petunia x hybrida* | Q43716 (UniProt) |

